# Social mobility and biological aging among older adults in the United States

**DOI:** 10.1101/2021.10.19.465042

**Authors:** GH Graf, Y Zhang, BW Domingue, KM Harris, M Kothari, D Kwon, P Muennig, DW Belsky

## Abstract

Lower socioeconomic status is associated with faster biological aging, the gradual and progressive decline in system integrity that accumulates with advancing age. Efforts to promote upward social mobility may therefore extend healthy lifespan. However, recent studies suggest that upward mobility may also have biological costs related to the stresses of crossing social boundaries. We analyzed blood-chemistry and DNA methylation (DNAm) data from n=9286 participants in the 2016 Health and Retirement Study (HRS) Venous Blood Study to test associations of life-course social mobility with biological aging. We quantified social mobility from childhood to later-life using data on childhood family characteristics, educational attainment, and wealth accumulation. We quantified biological aging using three DNA methylation “clocks” and three blood-chemistry algorithms. We observed substantial social mobility among study participants. Those who achieved upward mobility exhibited less-advanced and slower biological aging. Associations of upward mobility with less-advanced and slower aging were consistent for blood-chemistry and DNAm measures of biological aging and were similar for men and women and for Black and White Americans (Pearson-r effect-sizes ~0.2 for blood-chemistry measures and the DNAm GrimAge clock and DunedinPoAm pace-of-aging measures; effect-sizes were smaller for the DNAm PhenoAge clock). Analysis restricted to educational mobility revealed differential effects by racial identity, suggesting that mediating links between educational mobility and healthy aging may be disrupted by structural racism. In contrast, mobility producing accumulation of wealth appeared to benefit White and Black Americans equally, suggesting economic intervention to reduce wealth inequality may have potential to heal disparities in healthy aging.

**Significance Statement:** Upward social mobility may disrupt effects of early-life disadvantage on aging-related health decline. However, the stresses of crossing social boundaries can have biological costs. To investigate the balance of these forces, we analyzed social mobility from reports of childhood circumstances, education, and later-life wealth in 9,286 older adults in the US Health and Retirement Study. We quantified life-course health impacts of social mobility from blood-chemistry and DNA-methylation analysis of biological aging. We found that educational mobility alone benefited Black Americans less than White Americans, whereas mobility that produced accumulation of wealth into later-life was associated with delayed biological aging across social categories. Black-White disparities in healthy-aging outcomes of educational mobility may reflect inequalities in social gains realized from education.

## INTRODUCTION

Children who grow up poor get sick and die younger than their peers who grow up in more socioeconomically-advantaged families (1, 2). This inequality is mediated by a range of chronic diseases and health problems that become more frequent as individuals age, suggesting that childhood disadvantage may actually accelerate the aging process (3). Breakthroughs in aging biology have revealed a set of molecular changes that accumulate as individuals grow older, undermining resilience and driving vulnerability to multiple different chronic diseases, disability, and mortality (4). While there is currently no gold standard to measure this progressive loss of system integrity, several methods have been proposed (5). Current state-of-the-art methods are algorithms that combine information from multiple clinical or genomic measurements to track changes that occur in peoples’ bodies as they age. In longitudinal studies that track children through midlife, these algorithm-based methods reveal that people who grew up in disadvantaged households are biologically older and are aging more rapidly as adults as compared to peers with more advantaged childhoods, and vice versa (6–10). In crosssectional studies, children and adults in higher socioeconomic-status households exhibit less-advanced and slower biological aging as compared to those with lower socioeconomic status. These findings suggest that upward socioeconomic mobility, in which children climb the social ladder to achieve higher levels of status attainment than their family of origin, may interrupt processes that accelerate aging.

Conversely, upward mobility may also have biological costs. The stresses of climbing the social ladder, such as prolonged, high-effort coping, can damage health (11–15). This effect may be especially pronounced for groups facing structural barriers to upward mobility, such as Black Americans. If upward mobility accelerates biological aging, then interventions to build opportunity for at-risk children will need to devise additional strategies to offset potential health costs.

We tested if life-course socioeconomic mobility was associated with slower or faster biological aging in a national sample of U.S. adults, the U.S. Health and Retirement Study. We quantified social mobility from childhood to later-life based on retrospective reports by participants about their childhood socioeconomic conditions and structured interviews about household wealth. We quantified biological aging using DNA methylation- and physiology-based methods. Our analysis proceeded in three steps. We first tested how life-course socioeconomic disadvantage was associated with accelerated biological aging. We next tested whether upward social mobility was associated with blunting or amplification of associations between early-life socioeconomic disadvantage and accelerated biological aging. Finally, we tested if associations of mobility with biological aging were consistent for men and women and for Black and White adults to evaluate the hypothesis that the cost of social mobility could be more pronounced for groups who face structural barriers to upward mobility. We conducted parallel analysis of participants’ educational mobility.

## METHODS

### Data and Participants

We analyzed data from participants in the 2016 Health and Retirement Study (HRS) who provided blood chemistry and DNA methylation data in the Venous Blood Study. The Health and Retirement Study (HRS) is a nationally representative longitudinal survey of U.S. residents ≥50 years of age and their spouses (https://hrs.isr.umich.edu/documentation). The HRS has been fielded every two years since 1992. A new cohort of 51-56 year-olds and their spouses is enrolled every six years to maintain representativeness of the U.S. population over 50 years of age. Participants are asked about four broad areas: income and wealth; health, cognition and use of healthcare services; work and retirement; and family connections. As of the 2016 data release, the HRS included data collected from 42,515 individuals in 26,600 households. The 2016 Venous Blood Study (VBS) collected biomarker data from a subset of HRS participants who consented to a venous blood draw, as part of a larger effort to understand biological mechanisms linking social factors and health (n=9286). DNA methylation assays were done on a non-random subsample of VBS participants representative of the larger HRS sample (n=3989). We linked HRS data curated by RAND Corporation with new data collected as part of HRS’s 2016 Venous Blood Study ((16, 17). Our final analytic sample included all individuals who 1) participated in the 2016 wave of the HRS, 2) provided biomarker and/or DNA methylation data through the VBS, and 3) provided retrospective reports of socioeconomic indicators in childhood, middle adulthood, and later-life. The final analytic sample was 9,255 for analyses using biomarker measures of biological aging and 3,976 for analyses using DNA methylation measures of biological aging. Comparison of Venous Blood Study participants to the full HRS is reported in **Supplemental Table S1 Panel A**.

### Measures

#### Biological Aging

Biological aging is the gradual and progressive decline in system integrity that occurs with advancing chronological age, mediating aging-related disease and disability (18). While there is no gold standard measure of biological aging (5), current state-of-the-art methods use machine learning to sift through large numbers of candidate biomarkers and parameterize algorithms that predict aging-related parameters, including chronological age, mortality risk, and rate of decline in system integrity. Algorithms are developed in reference datasets and can then be applied to new datasets to test hypotheses.

For our analysis, we selected three blood-chemistry measures and three DNA methylation measures of biological aging shown in previous work to predict morbidity and mortality (6, 19–23), and which also demonstrated more advanced or more rapid aging in low socioeconomic status adults (6, 7, 24, 25). We compared different measures of biological aging to evaluate robustness of findings and to compare the sensitivity of blood-chemistry and DNA methylation biological-aging algorithms.

Blood-chemistry measures of biological aging were derived using three published methods: Phenotypic Age (20), Klemera-Doubal Method (KDM) Biological Age (26), and Homeostatic Dysregulation (27) applied to clinical chemistries and complete blood count data from venous blood draws. Algorithm parameterization for the KDM Biological Age and Homeostatic Dysregulation measures was conducted using the NHANES III data. PhenoAge parameterization was taken directly from the published article by Levine et al. (20). All blood chemistry measures were implemented in the HRS data using the BioAge R package (https://rdrr.io/github/dayoonkwon/BioAge/) (28).

DNA methylation measures of biological aging were obtained from the HRS (16). We conducted analysis of three measures: the PhenoAge clock (20), the GrimAge clock (29), and the DunedinPoAm Pace of Aging (6).

We refer to individual differences in the measures of biological aging as reflecting more/less advanced biological aging in the case of the blood-chemistry measures and DNA methylation clocks and as reflecting faster/slower aging in the case of the DunedinPoAm DNA methylation measure. The blood-chemistry measures and the DNA-methylation clocks have similar interpretation: They quantify how much biological aging a person has experienced up to the time of measurement. For those whose clock-ages are older/younger than their chronological ages, biological aging is more/less “advanced” relative to expectation. In contrast, DunedinPoAm measures how rapidly a person has been aging over the recent past. Values above the benchmark rage of 1 year of change per 12-month interval indicate “faster” biological aging, whereas values below 1 indicate “slower” biological aging. Measures are described in more detail in **Table 1** and **Supplemental Methods Section I**.

**Table 1.** Measures of Biological Aging Included in Analysis. The table reports the six measures of biological aging included in analysis. For each measure, the table reports the criterion used to develop the measure and the interpretation of the measure’s values. Criterion refers to the quantity the biological aging algorithm was developed to predict. Interpretation refers to the inference about biological aging that can be made on the basis of the values of the measure.

| Measure |  | Criterion | Interpretation |
| --- | --- | --- | --- |
| **Blood-Chemistry Measures.** All algorithms were parameterized using data from NHANES III and included the following blood chemistries: albumin, alkaline phosphatase, creatinine, C-reactive protein (log), white blood cell count, lymphocyte %, mean cell volume, and red cell distribution width. PhenoAge additionally included glucose. KDM Biological Age and Homeostatic Dysregulation additionally included glycated hemoglobin (HbA1C). PhenoAge and KDM Biological Age algorithms included information about chronological age. For analysis, PhenoAge and KDM Biological Age were differenced from chronological age to calculate biological-age advancement values. |  |  |  |
|  | PhenoAge | Mortality | Age at which the participant's biomarker-predicted mortality risk matches the norm in the reference population (NHANES III). |
|  | KDM Biological Age | Chronological Age | Age at which the participant's biomarker-predicted physiological integrity matches the norm in the reference population (NHANES III). |
|  | Homeostatic Dysregulation | Deviation from healthy youth | Log biomarker-Mahalanobis distance of participant from young, healthy reference population (non-obese NHANES III participants aged 20-30 years). |
| **DNA-Methylation Measures.** DNA-methylation measures were developed from analysis of genome-wide DNA methylation measured on Illumina 27k and 450k arrays in a range of different datasets. The Horvath Clock was developed from analysis of 82 different datasets. The Hannum Clock was developed from analysis of research volunteers at UC San Diego, University of Southern California, and West China Hospital. The PhenoAge Clock was developed from analysis of NHANES III and the InCHIANTI Study. The GrimAge clock was developed from analysis of the Framingham Heart Study Offspring Cohort. The DunedinPoAm Pace of Aging was developed from analysis of the Dunedin Study. DNA methylation measures were calculated by the HRS investigators. For analysis, DNA methylation clocks were residualized on chronological age to calculate biological-age advancement values. |  |  |  |
| **Second Generation DNA Methylation Clocks** |  |  |  |
|  | PhenoAge Clock | Blood-chemistry PhenoAge | DNAm prediction of the age at which the participant's biomarker-predicted mortality risk matches the norm in the NHANES III reference population (based on analysis of the InCHIANTI Study). |
|  | GrimAge Clock | Mortality | Age at which the participants' DNAm-predicted mortality risk matches the norm in the reference population (Framingham Heart Study Offspring cohort). The GrimAge clock was derived by first developing DNAm surrogates for blood proteins and smoking history and then developing a mortality prediction model based on these DNAm surrogates, sex, and chronological age. |
| **Pace of Aging** |  |  |  |
|  | DunedinPoAm Pace of Aging | Change over 12-years of follow-up in 18 system-integrity biomarkers | Years of physiological decline experienced per 1 year of calendar time over the recent past. DunedinPoAm was developed by modeling a composite of change scores for 18 biomarkers of organ system integrity from DNAm data. The expected value of DunedinPoAm in midlife adults is 1. Values >1 indicate accelerated aging. Values <1 indicate slowed aging. |

#### Social Mobility

We measured social mobility from participant reports about their socioeconomic circumstances before age 16, and from structured interviews about later-life wealth conducted by HRS between 1993 and 2016.

#### Childhood Social Origins

We constructed a childhood social origins index based on participants’ retrospective reports about their family’s general financial circumstances relative to other families, their father’s occupation, the family’s experiences of economic hardship, and their parents’ educational attainment. We composed the childhood social origins index as follows: First, we conducted principal components analysis of financial circumstances, father’s occupation, financial hardship, and parents’ education scores for HRS participants with complete data on all items (n=30,062). Second, we imputed missing values for father’s occupation and parents’ education set to the means for groups of participants matched on race, HRS birth cohort, and family financial circumstances score. Third, we applied loadings from the complete-case principal components analysis to compute factor scores for all participants. For analysis, we converted factor scores to Z scores (M=0, SD=1) and percentile ranks within 5-year birth cohorts. For the final childhood social origins index, higher values indicate more advantaged families of origin and lower values indicate less advantaged families of origin.

#### Later-life Socioeconomic Attainment

We measured later-life socioeconomic attainment from wealth data collected during structured interviews with participants about assets and liabilities over the course of multiple waves of participation in the HRS. Wealth data were chosen on the basis of evidence that wealth is more informative about social status in older adults as compared with income and educational level (30, 31) and shows clear associations with a range of aging-related health and functional deficits (32). We used wealth data compiled by RAND Corporation (33) and merged with data distributed by HRS. Because wealth data were combined across multiple years of measurement, we inflated all values to constant 2012 dollars. We applied an inverse-hyperbolic-sine transformation to reduce skew (34). Finally, we applied a theta transformation including adjustment for age and sex to achieve an approximately normal distribution of values (35). For analysis, we converted the transformed wealth values to Z scores (M=0, SD=1) and percentile ranks to form later-life socioeconomic attainment scores. Higher values of the later-life socioeconomic attainment score indicate higher levels of attainment and lower values indicate lower levels of attainment.

#### Mobility

We measured social mobility from childhood to later-life using two complementary approaches. (1) Residualized-change: we regressed participants’ later-life-socioeconomic-attainment z-score on their childhood-social-origins z-score and calculated residual values as a measure of mobility. (2) Difference-score: we calculated mobility as the difference between the later-life socioeconomic attainment z-score and the childhood social origins z-score. These two measures of mobility were highly correlated (r=.76). We conducted parallel analysis of both measures. We also conducted analysis of social mobility measured in terms of percentile-rank change from childhood to later-life using both residualized-change and difference score approaches. Details of social mobility variables are reported in **Supplementary Table S1, Panel B**.

#### Educational Mobility

We conducted parallel analysis of mobility from participant reports about their own education and the education of their parents.

#### Parental Education

We coded parental education in three categories based on years of schooling. To account for secular trends in educational attainment, we normalized parental educational attainments to five-year birth cohorts of participants. We classified those with educational attainment below the 25^th^ percentile as having low educational attainment, those with educational attainment between the 25^th^ and 75^th^ percentile as having average educational attainment, and those with educational attainment above the 75^th^ percentile as having high educational attainment. We assigned the highest attainment category of either parent as the participant’s parental educational attainment. This approach classified 20% of participants with low parental educational attainment, 57% with average parental educational attainment, and 23% with high parental educational attainment.

#### Participant Education

We coded participant education into three categories: those who had not graduated from high school (22%), those who had graduated from high school but had not completed a college degree (53%), and those who had completed at least a college degree (25%).

#### Educational mobility

We calculated educational mobility as the difference in education categories between participants and their parents. We assigned index scores of 1, 2, and 3 to respondents’ educational attainment (less than high school, high school, more than high school) and their parents’ educational attainment (low, medium, high). We calculated educational mobility by subtracting parental education index scores from participant education index scores, such that negative values represent downward social mobility and positive values represent upward social mobility (range −2 to 2, mean=0.02, SD=0.71). Details of educational mobility variables are reported in **Supplementary Table S1, Panel C**.

### Analysis

We used linear regression to test associations of social mobility with biological aging using the following specification:

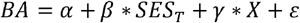

where *BA* is the measure of biological aging, *SES* is the socioeconomic circumstances measure (childhood social origins, later-life socioeconomic attainment, or social mobility), and *X* is a matrix of covariates. All models included covariate adjustment for chronological age, specified as a quadratic term, sex, whether the respondent self-identified as Hispanic, self-identified race (White, Black, Other), and the interactions of age terms with sex, race, and Hispanic ethnicity, *ε* represents the error term. The coefficient *β* tests the association of the SES measure with biological aging. We report results for z-score transformations of mobility in the main text and report results for both metrics in the Supplemental Tables.

We tested if associations of social mobility with biological aging varied by childhood socioeconomic status, sex, or race by adding cross-product interaction terms to our models:

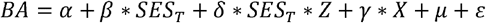

Where *BA*, *SES*, and *X* terms are the same as in the previous model and *Z* denotes the stratification variable (childhood socioeconomic position, sex, or Black/White racial identity). The coefficient *δ* tests the hypothesis that the association of mobility with biological aging varies across levels/strata of *Z*.

We used the same models to test associations of educational mobility with biological aging. In these models, the SES terms were replaced with terms for parents’ educational attainment, participants’ educational attainment, and the difference in attainments between parents and participants.

For all models, effect-sizes are scaled in standard deviation units of the outcome measure. Positive effect-sizes indicate more-advanced or faster biological aging; negative effect-sizes indicate less-advanced or slower biological aging. For social-mobility models, effectsizes are reported for a 1 standard deviation difference in the exposure. For educational mobility models, effect-sizes are reported for a single-category increases in educational attainment.

To account for non-independence of observations of couples who share a household, we clustered standard errors at the household level. We conducted all analyses in RStudio Version 1.3.1093.

## RESULTS

HRS participants included in analysis showed substantial social mobility (percentile-rank mobility SD=25). Compared to the full 2016 HRS sample, participants in the VBS subsample and the DNA methylation subsample for whom biological aging measures could be computed were somewhat more likely to be White and to experience more upward social mobility. Comparison of socio-demographic characteristics of the analysis sample to the full 2016 HRS panel is reported in **Supplemental Table S1** and **Supplemental Figure S5**.

### HRS participants who grew up in more socioeconomically advantaged households exhibited less-advanced and slower biological aging in later-life

We combined participants’ retrospective reports about their parents’ education, childhood experiences of economic hardship, and perceptions of their family’s relative socioeconomic standing into a single index of childhood social origins. Participants who grew up in more socioeconomically advantaged households exhibited less-advanced and slower biological aging across all six aging measures included in our analysis (effect-size range *β*=[-0.07,-0.03], where ‘*β*’ represents an effect-size denominated in standard-deviations of biological aging per standard-deviation difference in social origins; **Supplemental Figure S1, Supplemental Table S2)**. However, effect-sizes were small, consistent with a prior report from this cohort (19).

### HRS participants with higher levels of later-life socioeconomic attainment exhibited less-advanced and slower biological aging

We measured later-life socioeconomic attainment from household wealth data collected from structured interviews with participants about their assets and liabilities and compiled by RAND corporation. Participants with higher levels of attainment exhibited less-advanced and slower biological aging across all six measures of biological aging included in our analysis (attainment Z-score range *β*=[-0.25,-0.18], except for DNAm PhenoAge (*β*=-0.09), where ‘*β*’ represents an effect-size denominated in standard-deviations of biological aging per standard-deviation difference in attainment; **Supplemental Figure S1, Supplemental Table S2)**. These effect-sizes were larger relative to the association of childhood social origins with biological aging.

### HRS participants who climbed up the social ladder showed less-advanced and slower biological aging in later-life

We measured socioeconomic mobility in two ways. First, we computed mobility as the difference in the level of later-life socioeconomic attainment achieved from the level of attainment expected based on childhood social origins (the residual from a regression of later-life socioeconomic attainment on childhood social origins; hereafter “residualized-change mobility”). Participants with more upward mobility exhibited less-advanced and slower biological aging (residualized-change mobility Z-score range *β*=[-0.23,-0.16], except for DNAm PhenoAge (*β*=-0.09), where ‘*β*’ represents an effect-size denominated in standard-deviations of biological aging per standard-deviation difference in mobility). Second, we computed mobility as a simple difference score (later-life socioeconomic attainment-childhood social origins; hereafter “delta mobility”). Consistent with results from our first approach, participants with more upward mobility exhibited less-advanced and slower biological aging (delta mobility Z-score range *β*=[-0.09,-0.06] except for DNAm PhenoAge (*β*=-0.02)); **Figure 2, Supplemental Figure S1, Supplemental Table S2a)**.

**Figure 1.**
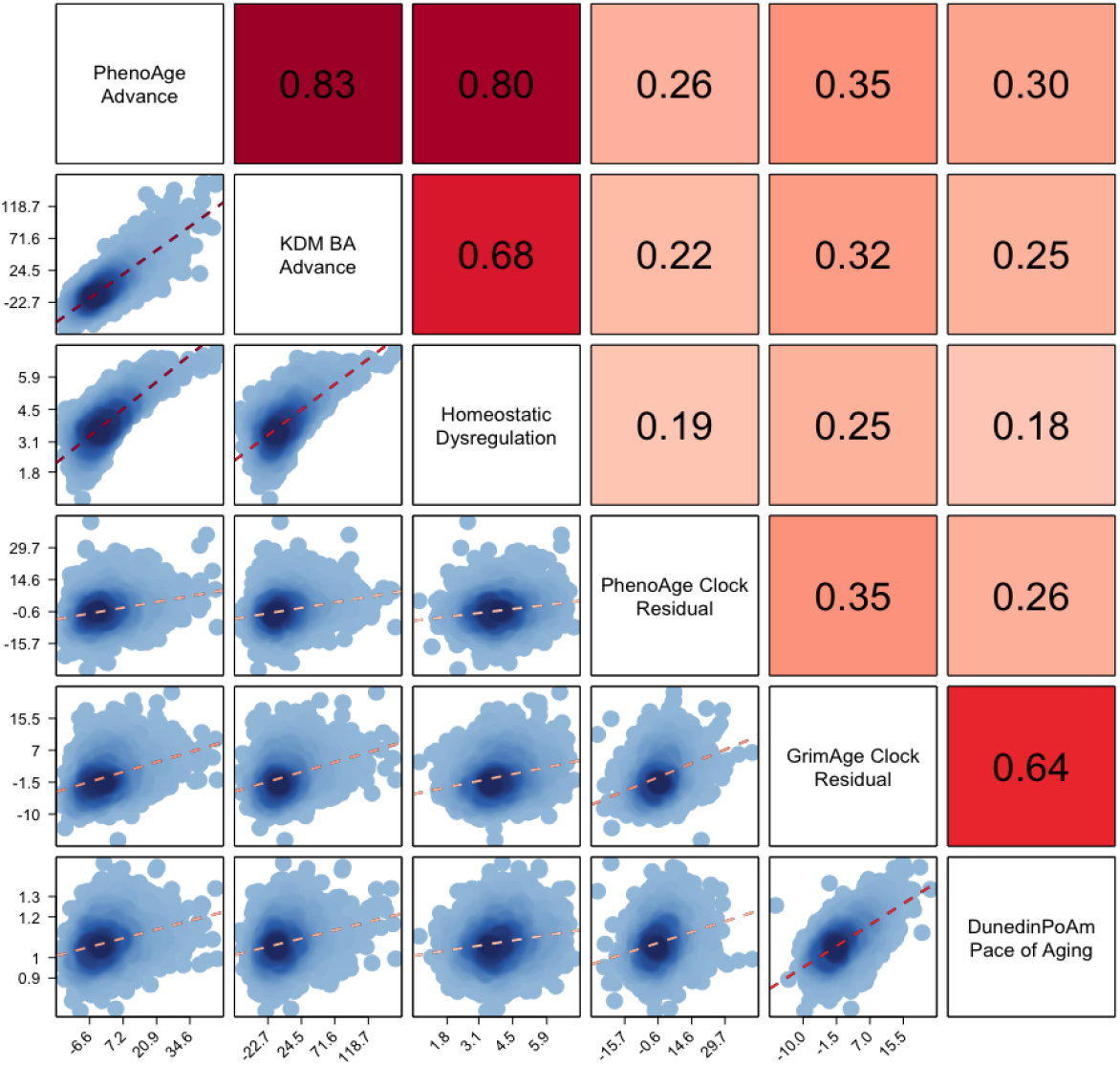
Correlations among three blood-chemistry and three DNA-methylation measures of biological aging among Black and White participants in the US Health and Retirement Study. Biological aging measure labels are listed on the matrix diagonal. Pearson correlations are shown above the diagonal. Correlations are reported for the biological aging measures listed below and to the left of the cell. Scatterplots and linear fits illustrating associations are shown below the diagonal. The Y axis of the plots corresponds to the biological aging measure to the right of the cell. The X axis of the plots corresponds to the biological aging measure above the cell. Sample sizes for correlations among blood-chemistry measures are n=9255. Sample sizes for correlations between blood-chemistry and DNA-methylation measures and among DNA-methylation measures are n=3976.

**Figure 2:**
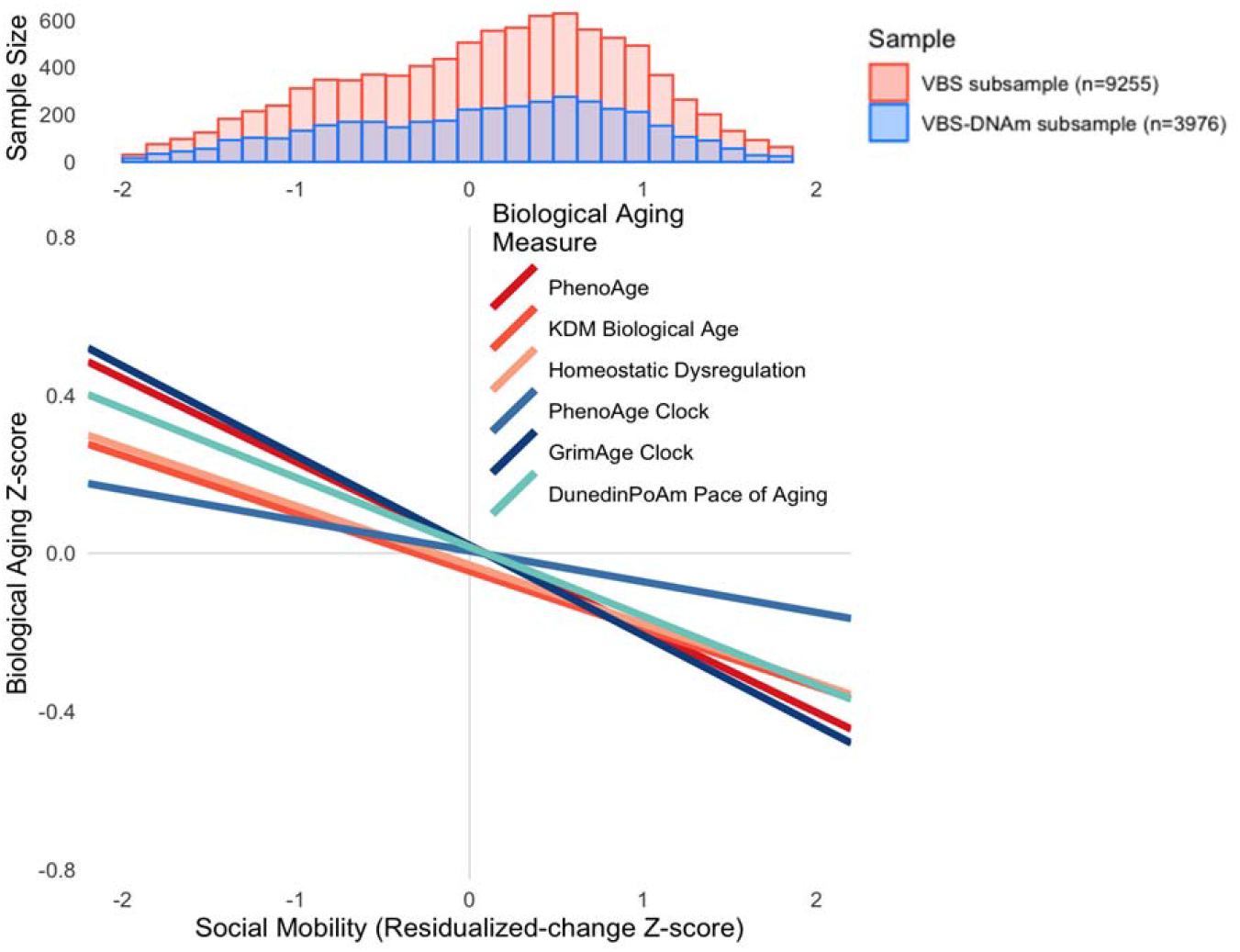
Effect-sizes for associations of life course social mobility with three blood-chemistry and three DNA methylation measures of biological aging. The histogram at the top of the figure shows the distribution of social mobility in percentile rank terms in the full HRS Venous Blood Study (n=9255; red bars) and the DNA methylation subsample (n=3976; blue bars). The line plot at the bottom of the figure shows the association of social mobility with six measures of biological aging. Blood-chemistry-based measures are plotted in red shades. DNA methylation measures are plotted in blue shades. The figure shows that, across methods, upward social mobility was associated with less-advanced and slower biological aging.

We conducted sensitivity analyses to evaluate consistency of associations between social mobility and biological aging across three sets of groups facing different barriers to social mobility: those who grew up in more as compared with less disadvantaged families; women as compared with men; and Black as compared with White Americans.

#### Childhood social origins

The association between upward social mobility and biological aging was similar across the distribution of childhood social origins (interaction p-values>0.237). This finding remained consistent when we restricted analysis to participants in the middle 50% of the childhood social origins distribution. Results are reported in **Supplemental Table S3** and **Supplemental Figure S2**.

#### Sex

For both women and men, upward social mobility was associated with less-advanced and slower biological aging (for women, residualized-change mobility effect-size range *β*=[-0.26,-0.15] except for DNAm PhenoAge (*β*=-0.08), delta mobility effect-size range *β*=[-0.10,-0.05] except for DNAm PhenoAge (*β*=-0.004); for men, residualized-change mobility effect-size range *β*=[-0.28,-0.12] except for DNAm PhenoAge (*β*=-0.07), delta mobility effectsize range *β*=[-0.10,-0.04]). In residualized-change analysis, effect-sizes for blood-chemistry PhenoAge and Homeostatic Dysregulation measures of biological aging indicated somewhat stronger associations of mobility with biological aging for women as compared to men (interaction term range *β*=[-0.09,-0.04]. However, DNA methylation measures of aging did not show consistent differences, and effect-size differences were not generally statistically significant at the alpha=0.05 level. In delta-mobility analysis, effect-size differences between men and women were not statistically significant at the alpha=0.05 level (p>0.113). Results are reported in **Supplemental Table S4a** and **Supplemental Figure S3**.

#### Racial identity

For both White and Black adults, upward social mobility was associated with less-advanced and slower biological aging (for Black adults, residualized-change mobility effect-size range *β*=[-0.25,-0.16] except for DNAm PhenoAge (*β*=-0.09), delta mobility effectsize range *β*=[-0.11,-0.08] except for DNAm PhenoAge (*β*=-0.05); for White adults, residualized-change mobility effect-size range *β*=[-0.25,-0.15] except for DNAm PhenoAge (*β*=-0.09), delta mobility effect-size range *β*=[-0.09,-0.03)). Effect-size differences between White and Black adults were not statistically significant at the alpha=0.05 level (p-values for tests of interactions.052). Results are reported in **Supplemental Table S4b** and **Supplemental Figure S4)**.

The consistency of effect-sizes for social-mobility associations with biological aging between White and Black HRS participants contrasts with reports that associations of socioeconomic attainment with health may be weaker for Black as compared to White Americans (12, 14, 15, 36). In these studies, socioeconomic attainment was measured from education. We therefore repeated our analysis with a mobility measure derived by comparing educational attainments of participants to those of their parents (hereafter “educational mobility”).

#### Analysis of educational mobility

Effect-sizes for educational-mobility associations with biological aging were somewhat smaller than effect-sizes for social-mobility associations (range *β*=[-0.13,-0.02], **Supplemental Table S2b).** As in analysis of social mobility, blood-chemistry measures of biological aging indicated somewhat larger effect-sizes for women as compared to men (for women, effect-size range *β*=[-0.14,-0.06]; for men, effect-size range *β*=[-0.13,0.03]; **Supplemental Table S4c)**. For Black and White adults, upward educational mobility was associated with less-advanced and slower biological aging (for Black adults, effect-size range *β*=[-0.20,-0.04]; for White adults, effect-size range *β*=[-0.17,-0.05]). Effect-sizes were smaller in Black as compared to White adults with the exception of DunedinPoAm analysis, which showed the reverse pattern. However, differences were not statistically significant at the alpha=0.05 level in the DNA methylation sample. Results are reported in **Supplemental Table S4d**.

## DISCUSSION

We tested how life-course socioeconomic mobility related to healthy aging in a national sample of older adults in the United States. We measured healthy aging using blood-chemistry and DNA-methylation measures of the state and pace of biological aging. There were three main findings. First, older adults who had grown up in socioeconomically at-risk families and those who had accumulated less wealth across their lives exhibited more-advanced and faster-paced biological aging as compared to those who grew up in more socioeconomically-advantaged families. Second, those who overcame early-life disadvantage and climbed the social ladder to achieve upward mobility had less-advanced and slower-paced biological aging in later life as compared with those who were non-mobile or downwardly mobile. Third, upward-mobility associations with healthy aging were generally consistent for men and women, for White and Black adults, and for those who started life at different levels of socioeconomic position. In sum, we did not find evidence of net biological costs associated with the stresses of climbing the social ladder. Instead, findings suggest that upward socioeconomic mobility contributes to healthy aging, including in groups that face structural barriers to socioeconomic attainment.

Our findings were consistent across metrics of aging derived from different biological levels of analysis and developed using different models of the aging process. Childhood socioeconomic disadvantage, lower levels of wealth in later-life, and downward social mobility were each associated with more-advanced/faster biological aging across three blood-chemistry measures (blood-chemistry PhenoAge, KDM Biological Age, and Homeostatic Dysregulation) and three DNA methylation measures (PhenoAge Clock, GrimAge Clock, and DunedinPoAm Pace of Aging), although effect-sizes were smaller for the DNA-methylation PhenoAge Clock. These six measures comprise biological clocks that estimate the extent of aging in a person (KDM Biological Age, the PhenoAge measures, and GrimAge), a measure of physiologic deviation from a healthy, youthful state (homeostatic dysregulation), and a Pace of Aging measure that estimates the ongoing rate of decline in system integrity (DunedinPoAm). Consistency of findings across biological levels of analysis and conceptually distinct measures of aging builds confidence in the robustness of the association of upward social mobility with healthy aging.

Our results contrast with some previous reports suggesting that there may be physical health costs from upward social mobility (12–15, 36). A possible explanation is that we measured life-course socioeconomic attainment from data on wealth accumulation whereas previous studies had focused on educational attainment (12, 14, 15, 36). When we conducted analysis of educational mobility, our findings were more consistent with prior studies; effectsizes for upward educational mobility were 2-4 times larger in analysis of White as compared to Black participants, with the exception of the DunedinPoAm Pace of Aging measure, for which the educational-mobility effect-size was larger in Black as compared to White participants. (In tests of interaction, effect-size differences were not statistically significant at the alpha=0.05 level for any of the measures.)

The difference in findings in analysis of social mobility as compared to educational mobility may reflect differences in the life stage timing of the measurements used to quantify these processes and in the ways that the different mobility processes themselves affect the lives of Black and White Americans. The data we used to quantify life-course attainment in social mobility analysis was derived from structured interviews the HRS conducted with participants about their assets and liabilities during follow-ups spanning 1992-2016. These data capture levels of resources participants accumulated across their lives and had access to during the years leading up to the blood draws from which we derived our measures of healthy aging. Conversely, participants mostly completed their education decades before aging measurements were taken. Educational attainment plausibly represents young adult potential to accumulate socioeconomic and material resources that may affect healthy aging. However, this potential is likely unequally realized for Black and White Americans (37). One explanation for why educational mobility showed weaker associations with healthy aging in Black as compared to White participants is that Black Americans, who face racism in educational, work, and community environments, and who are part of extended family networks with lower levels of resources overall, don’t realize the same social and material gains from their education as their White peers, e.g. (38, 39).

We acknowledge limitations. There is no gold standard measure of biological aging (5). Our conclusions are circumscribed by the precision and validity of available measurements. Our analysis included DNA-methylation- and blood-chemistry-based measures. Other proposed levels of analysis for quantification of biological aging include proteomics, metabolomics, and physical performance tests. Ultimately, integrating information across levels of analysis may yield more precise measurements (40). However, consistency of results across different blood-chemistry and DNA-methylation methods builds confidence in findings. Social mobility was measured from participant-reported information. Reporting biases cannot be ruled out. Childhood socioeconomic circumstances, which were retrospectively reported, may be subject to recall bias. If aging trajectories affect recall of early-life adversity, or if participants’ anchoring their responses to different perceptions of normative socioeconomic conditions is related to other causes of aging, our findings may over- or underestimate the true effects of social mobility on healthy aging. Studies are needed that can link measures of biological aging with administrative records that objectively record dimensions of social mobility. Our sample was made up of adults aged 50 years and older and their spouses. To the extent that socioeconomic disadvantage and downward mobility are associated with premature mortality, our sample may underrepresent the most at-risk population segments, potentially biasing our results towards the null. Further, mortality differences across demographic groups mean that differences between Black and White participants, and between men and women, may be underestimated. Participation biases may compound this survival bias, especially for Black-White comparisons; Black participants in the Venous Blood Study were younger and healthier than the full sample of Black participants in the HRS (41). Our estimates of Black-White disparities are therefore likely to be conservative.

The observation that upward social mobility is associated with slower biological aging builds on evidence that people with more socioeconomic resources appear biologically younger than peers of the same chronological age with fewer socioeconomic resources (42). Mobility findings advance evidence for the hypothesis that intervention to promote economic well-being in adulthood can help to address disparities in healthy aging. But whether associations of upward mobility with slowed biological aging reflect effects of the resources acquired through upward mobility or from resources and characteristics that made mobility possible remains to be determined. A critical next step is to clarify when in the life course intervention can be most impactful and what mechanisms are most effective in delivering not just economic justice, but aging health equity. Collection of bio-samples from participants in studies of interventions to promote successful early-childhood development (43), increase educational attainment (44), and reduce poverty and promote stable housing and employment in adults (45, 46), can advance understanding of when and how interventions to address inequalities in social determinants of health can most powerfully affect inequalities in healthy aging.

## Acknowledgements

This research was supported by National Institute on Aging Grants R01AG061378 and R01AG066887, Russell Sage Foundation Grant 1810-08987, and the Jacobs Foundation. DWB is a fellow of the CIFAR CBD Network.

## Supplemental Methods

### Section I. Measures of Biological Aging

We selected three blood-chemistry measures of biological aging: Phenotypic Age (Levine et al., 2018), Klemera-Doubal Method (KDM) Biological Age (Klemera and Doubal, 2006), and Homeostatic Dysregulation (Cohen et al., 2013). The Phenotypic Age measure was developed from analysis of mortality risk; it represents the age at which a person’s blood-chemistry-predicted mortality risk would be approximately normal in the general population. The KDM Biological Age measure was developed from analysis of chronological age; it measures the age at which a person’s physiology would match the population norm. The Homeostatic Dysregulation measure was developed from analysis of deviation from a young, healthy reference sample; it quantifies how deviant a person’s physiology is from this reference.

We selected three DNA methylation measures of biological aging: the PhenoAge clock (Levine et al., 2018), the GrimAge clock (Lu et al., 2019), and the DunedinPoAm Pace of Aging measure (Belsky et al., 2020). The PhenoAge clock was developed from machine-learning analysis of the Phenotypic Age blood-chemistry measure. The GrimAge clock was developed in a two-stage analysis that first developed DNA methylation biomarkers of blood proteins and tobacco exposure and then fitted these DNA methylation biomarkers to mortality risk. Both the PhenoAge and GrimAge clocks measure the age at which a person’s mortality risk would be approximately normal in the population. DunedinPoAm was developed in a two-stage analysis that first developed a composite phenotype of within-person change over time in 18 biomarkers of organ system integrity, termed “Pace of Aging “, and then fitted DNA methylation data to predict that composite Pace of Aging.

## Supplemental Tables and Figures

**Supplemental Table S1.**
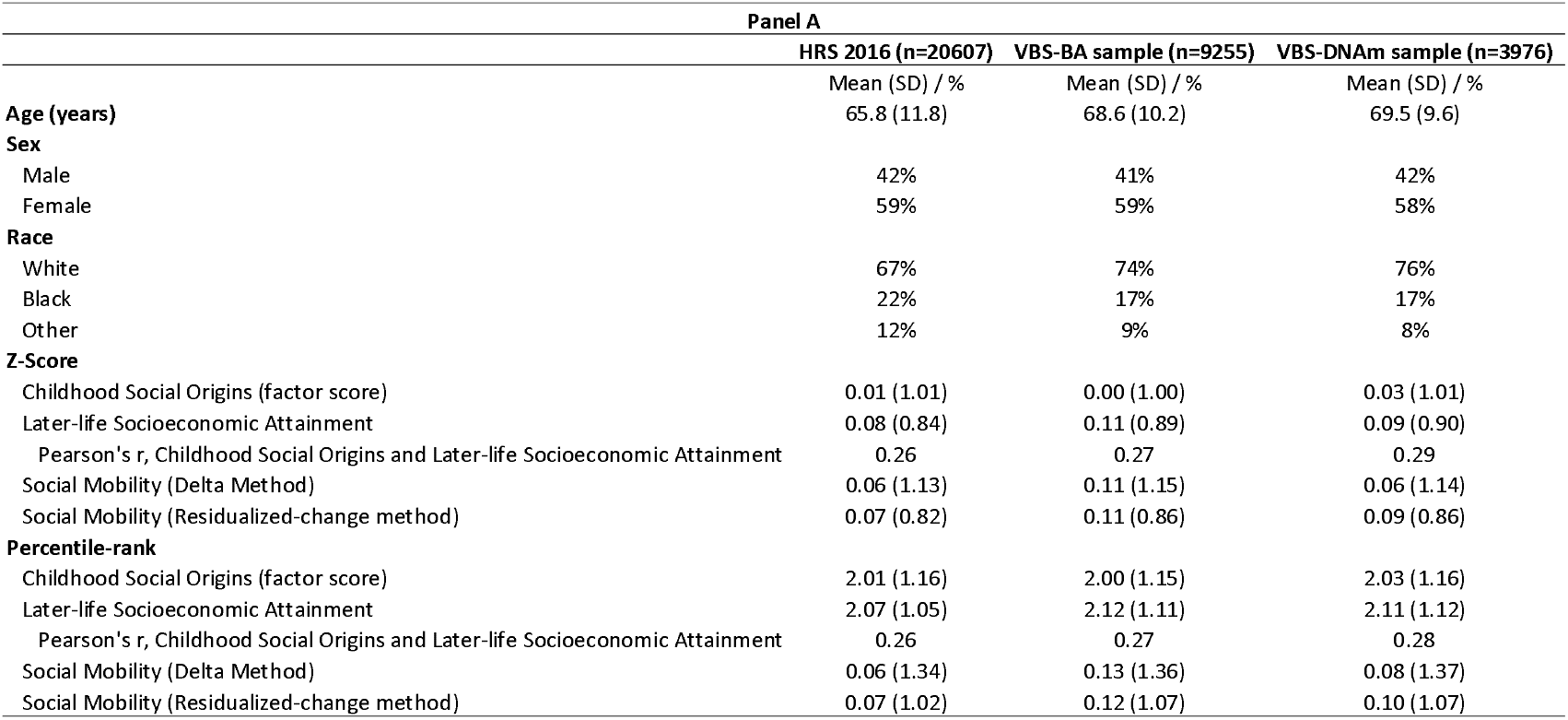
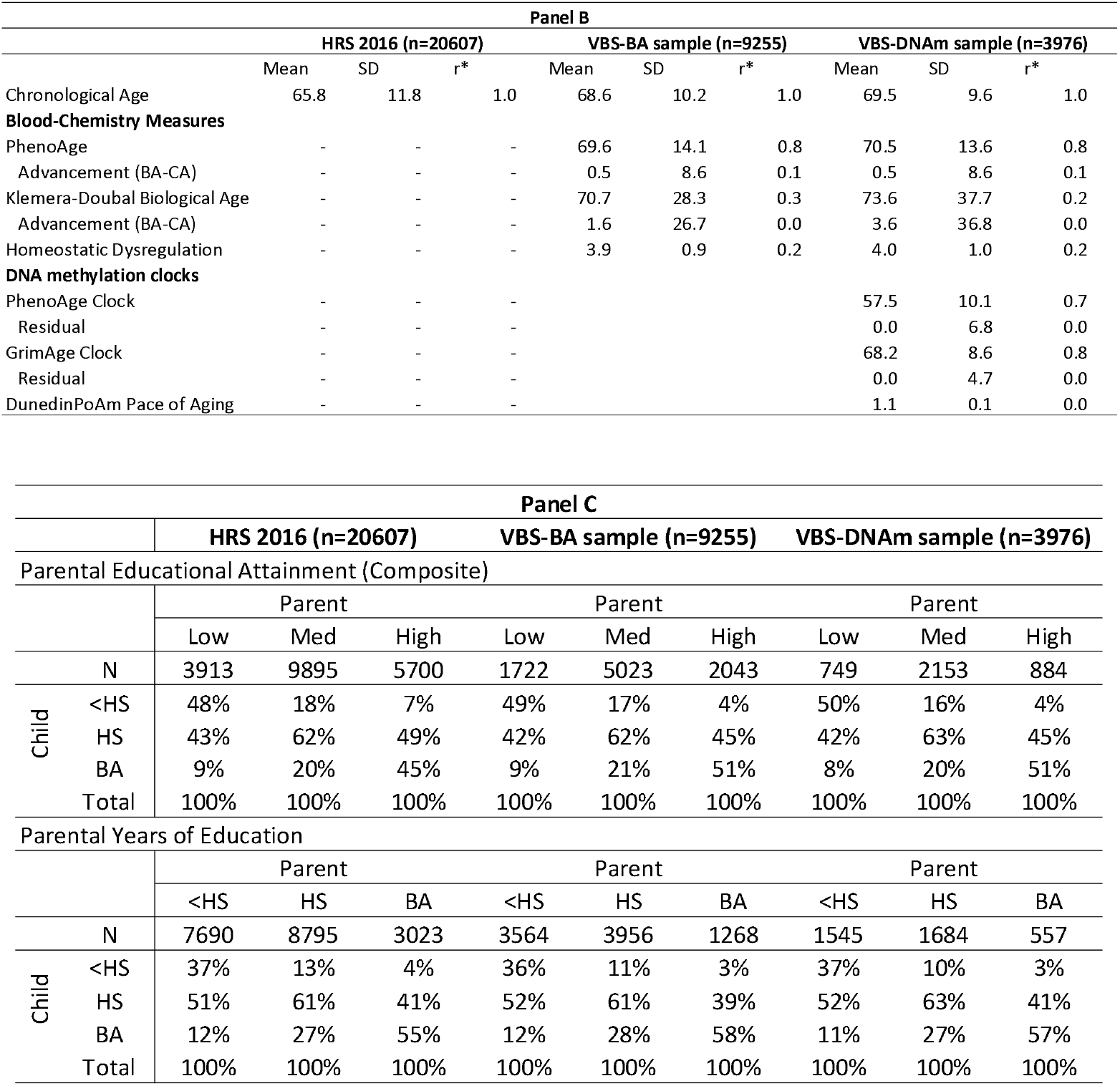
Comparison of demographic characteristics for participants in the 2016 wave of the US Health and Retirement Study and the subset of participants included in social mobility analysis. The full HRS sample consists of all participants in the 2016 Health and Retirement Study who provided demographic data and information on childhood socioeconomic status and household wealth (n=20607). The VBS-BA sample consists of all participants from the full HRS sample for whom biological-age values could be computed based on biomarker data obtained through the Venous Blood Study (n=9255). The VBS-DNAm sample consists of all participants from the full HRS sample for whom biological-age values could be computed based on biomarker data and DNA methylation data obtained through participation in the Venous Blood Study (n=3976). In Panel A, mean values for household wealth were calculated by first inflating wealth values to 2012 dollars using the Consumer Price Index and then taking the average across all HRS measurement waves. For analysis of mobility, wealth values were transformed according to the procedure described in the Methods section and converted to either Z-scores or percentile ranks. Residualized-change mean values reported in the table are not precisely equal to 1 for any of the samples because the regression to compute residuals included all HRS participants ever providing data on social origins and attainments (N=37,722). In Panel B, biological-age advancements were calculated by subtracting chronological age from biological age (BA-CA) for blood-chemistry measures. Age residuals were calculated by fitting a regression of biological age on chronological age to the full VBS-DNA Methylation Sample, then subtracting the fitted value from that estimated using DNA methylation clock calculations.

**Supplemental Table S2a.** Effect-sizes for associations of social origins, socioeconomic attainment, and social mobility with blood-chemistry and DNA-methylation measures of biological aging. The table reports effectsizes for associations of childhood socioeconomic status (SES), adult attainment (household wealth), and social mobility with blood-chemistry and DNA methylation measures of biological aging. For Z-score measures, effect-sizes are denominated in standard deviation units of biological age advancement per standarddeviation increment in the predictor. For percentile-rank measures, effect-sizes are denominated in standarddeviation units of biological age advancement per 25-percentile-rank increments of the predictor.

| **Panel I: Z-score Scaled Social Mobility Measures** |  |  |  |  |  |  |
| --- | --- | --- | --- | --- | --- | --- |
|  | VBS sample (n=9255) |  |  | VBS-DNAm sample (n=3976) |  |  |
|  | Estimate | CI | p | Estimate | CI | p |
| **Childhood Social Origins** |  |  |  |  |  |  |
| <u>Blood Chemistry Measures</u> |  |  |  |  |  |  |
| PhenoAge | -0.05 | [-0.07,-0.03] | **5.34E-06** | -0.05 | [-0.09,-0.02] | **8.97E-04** |
| Klemera-Doubal Biological Age | -0.04 | [-0.05,-0.02] | **1.21E-05** | -0.05 | [-0.08,-0.02] | **3.55E-03** |
| Homeostatic Dysregulation | -0.04 | [-0.06,-0.02] | **5.13E-05** | -0.05 | [-0.08,-0.02] | **1.74E-03** |
| <u>DNA Methylation Measures</u> |  |  |  |  |  |  |
| PhenoAge Clock | - | - | - | -0.03 | [-0.06,0.00] | 0.094 |
| GrimAge Clock | - | - | - | -0.07 | [-0.10,-0.04] | **1.32E-06** |
| DunedinPoAm | - | - | - | -0.04 | [-0.07,0.00] | **2.67E-02** |
| **Later-life Socioeconomic Attainment** |  |  |  |  |  |  |
| <u>Blood Chemistry Measures</u> |  |  |  |  |  |  |
| PhenoAge | -0.22 | [-0.25,-0.20] | **5.43E-72** | -0.24 | [-0.28,-0.20] | **1.24E-38** |
| Klemera-Doubal Biological Age | -0.15 | [-0.17,-0.13] | **1.65E-59** | -0.18 | [-0.21,-0.14] | **1.08E-20** |
| Homeostatic Dysregulation | -0.15 | [-0.18,-0.13] | **1.81E-43** | -0.19 | [-0.22,-0.15] | **7.24E-25** |
| <u>DNA Methylation Measures</u> |  |  |  |  |  |  |
| PhenoAge Clock | - | - | - | -0.09 | [-0.12,-0.05] | **7.04E-06** |
| GrimAge Clock | - | - | - | -0.25 | [-0.29,-0.22] | **5.82E-46** |
| DunedinPoAm | - | - | - | -0.19 | [-0.23,-0.15] | **2.14E-23** |
| **Social Mobility (Delta Method)** |  |  |  |  |  |  |
| <u>Blood Chemistry Measures</u> |  |  |  |  |  |  |
| PhenoAge | -0.08 | [-0.10,-0.06] | **5.19E-18** | -0.09 | [-0.11,-0.06] | **3.60E-10** |
| Klemera-Doubal Biological Age | -0.05 | [-0.07,-0.04] | **8.05E-15** | -0.06 | [-0.09,-0.03] | **6.75E-05** |
| Homeostatic Dysregulation | -0.05 | [-0.07,-0.04] | **3.89E-10** | -0.06 | [-0.09,-0.03] | **8.94E-06** |
| <u>DNA Methylation Measures</u> |  |  |  |  |  |  |
| PhenoAge Clock | - | - | - | -0.02 | [-0.05,0.00] | 0.081 |
| GrimAge Clock | - | - | - | -0.08 | [-0.10,-0.05] | **7.64E-10** |
| DunedinPoAm | - | - | - | -0.07 | [-0.10,-0.05] | **1.12E-07** |
| **Social Mobility (Residualized-change method)** |  |  |  |  |  |  |
| <u>Blood Chemistry Measures</u> |  |  |  |  |  |  |
| PhenoAge | -0.21 | [-0.23,-0.18] | **6.10E-62** | -0.23 | [-0.26,-0.19] | **2.18E-33** |
| Klemera-Doubal Biological Age | -0.14 | [-0.16,-0.12] | **5.20E-51** | -0.16 | [-0.20,-0.13] | **8.42E-17** |
| Homeostatic Dysregulation | -0.14 | [-0.17,-0.12] | **1.67E-36** | -0.18 | [-0.21,-0.14] | **8.68E-21** |
| <u>DNA Methylation Measures</u> |  |  |  |  |  |  |
| PhenoAge Clock | - | - | - | -0.08 | [-0.12,-0.04] | **5.88E-05** |
| GrimAge Clock | - | - | - | -0.23 | [-0.27,-0.20] | **1.16E-38** |
| DunedinPoAm | - | - | - | -0.18 | [-0.22,-0.14] | **3.25E-21** |

| **Panel II. Percentile-rank Scaled Social Mobility Measures** |  |  |  |  |  |  |
| --- | --- | --- | --- | --- | --- | --- |
| VBS sample (n=9286) |  |  |  | VBS-DNAm sample (n=3989) |  |  |
|  | Estimate | CI | p | Estimate | CI | p |
| **Childhood Social Origins** |  |  |  |  |  |  |
| <u>Blood Chemistry Measures</u> |  |  |  |  |  |  |
| PhenoAge | -0.04 | [-0.06,-0.02] | **1.50E-05** | -0.04 | [-0.07,-0.02] | **2.16E-03** |
| Klemera-Doubal Biological Age | -0.03 | [-0.04,-0.02] | **2.48E-05** | -0.04 | [-0.07,-0.01] | **6.41E-03** |
| Homeostatic Dysregulation | -0.03 | [-0.05,-0.02] | **1.34E-04** | -0.04 | [-0.07,-0.01] | **5.04E-03** |
| <u>DNA Methylation Measures</u> |  |  |  |  |  |  |
| PhenoAge Clock | - | - | - | -0.03 | [-0.05,0.00] | 0.073 |
| GrimAge Clock | - | - | - | -0.06 | [-0.09,-0.03] | **4.68E-06** |
| DunedinPoAm | - | - | - | -0.03 | [-0.06,0.00] | **4.02E-02** |
| **Later-life Socioeconomic Attainment** |  |  |  |  |  |  |
| <u>Blood Chemistry Measures</u> |  |  |  |  |  |  |
| PhenoAge | -0.18 | [-0.20,-0.16] | **9.96E-75** | -0.20 | [-0.23,-0.17] | **5.76E-41** |
| Klemera-Doubal Biological Age | -0.13 | [-0.14,-0.11] | **9.29E-62** | -0.15 | [-0.18,-0.12] | **9.78E-23** |
| Homeostatic Dysregulation | -0.13 | [-0.15,-0.11] | **8.82E-46** | -0.16 | [-0.19,-0.13] | **4.08E-27** |
| <u>DNA Methylation Measures</u> |  |  |  |  |  |  |
| PhenoAge Clock | - | - | - | -0.07 | [-0.10,-0.04] | **2.99E-06** |
| GrimAge Clock | - | - | - | -0.21 | [-0.24,-0.18] | **2.55E-48** |
| DunedinPoAm | - | - | - | -0.16 | [-0.19,-0.13] | **1.46E-24** |
| **Social Mobility (Delta Method)** |  |  |  |  |  |  |
| <u>Blood Chemistry Measures</u> |  |  |  |  |  |  |
| PhenoAge | -0.07 | [-0.09,-0.06] | **9.42E-23** | -0.08 | [-0.11,-0.06] | **4.71E-13** |
| Klemera-Doubal Biological Age | -0.05 | [-0.06,-0.04] | **2.84E-18** | -0.06 | [-0.08,-0.03] | **1.80E-06** |
| Homeostatic Dysregulation | -0.05 | [-0.06,-0.04] | **2.20E-13** | -0.06 | [-0.09,-0.04] | **3.29E-08** |
| <u>DNA Methylation Measures</u> |  |  |  |  |  |  |
| PhenoAge Clock | - | - | - | -0.02 | [-0.05,0.00] | 5.24E-02 |
| GrimAge Clock | - | - | - | -0.08 | [-0.10,-0.06] | **6.69E-13** |
| DunedinPoAm | - | - | - | -0.07 | [-0.09,-0.05] | **1.94E-09** |
| **Social Mobility (Residualized-change method)** |  |  |  |  |  |  |
| <u>Blood Chemistry Measures</u> |  |  |  |  |  |  |
| PhenoAge | -0.18 | [-0.20,-0.16] | **3.48E-66** | -0.19 | [-0.22,-0.16] | **1.24E-36** |
| Klemera-Doubal Biological Age | -0.12 | [-0.14,-0.10] | **3.02E-54** | -0.14 | [-0.17,-0.11] | **3.87E-19** |
| Homeostatic Dysregulation | -0.12 | [-0.14,-0.10] | **6.33E-40** | -0.15 | [-0.18,-0.12] | **6.95E-24** |
| <u>DNA Methylation Measures</u> |  |  |  |  |  |  |
| PhenoAge Clock | - | - | - | -0.07 | [-0.10,-0.04] | **2.69E-05** |
| GrimAge Clock | - | - | - | -0.20 | [-0.23,-0.17] | **7.31E-42** |
| DunedinPoAm | - | - | - | -0.15 | [-0.18,-0.12] | **9.10E-23** |

**Supplemental Table S2b.** Effect-sizes for associations of parental education, educational attainment, and educational mobility with blood-chemistry and DNA-methylation measures of biological aging. Effect-sizes are denominated in standard-deviation units of biological aging per one-category increase in educational attainment/ mobility. Categories of participant education are <high school, high school graduate, college graduate. Categories of educational social origins (parental education) are defined by the 25th and 75th percentiles of years of education completed by parents of participants grouped into 5-year birth cohorts. The final panel of the table shows results for parental education and educational mobility based on coding of parental education by the degree criteria used to code participant education.

|  | VBS-BA sample (n=9255) |  |  | VBS-DNAM sample (n=3976) |  |  |
| --- | --- | --- | --- | --- | --- | --- |
|  | Estimate | CI | p | Estimate | CI | p |
| **Parental education** |  |  |  |  |  |  |
| <u>Blood Chemistry Measures</u> |  |  |  |  |  |  |
| PhenoAge | -0.09 | [-0.12,-0.05] | **1.47E-06** | -0.10 | [-0.15,-0.05] | **2.23E-04** |
| Klemera-Doubal Biological Age | -0.05 | [-0.08,-0.03] | **1.26E-04** | -0.12 | [-0.18,-0.07] | **9.80E-06** |
| Homeostatic Dysregulation | -0.07 | [-0.11,-0.04] | **3.60E-06** | -0.11 | [-0.16,-0.05] | **6.67E-05** |
| <u>DNA Methylation Measures</u> |  |  |  |  |  |  |
| PhenoAge Clock | - | - | - | -0.09 | [-0.14,-0.04] | **7.90E-04** |
| GrimAge Clock | - | - | - | -0.17 | [-0.22,-0.12] | **9.31E-11** |
| DunedinPoAm | - | - | - | -0.06 | [-0.11,-0.01] | **2.93E-02** |
| **Participant Education** |  |  |  |  |  |  |
| <u>Blood Chemistry Measures</u> |  |  |  |  |  |  |
| PhenoAge | -0.18 | [-0.21,-0.15] | **6.04E-27** | -0.18 | [-0.23,-0.13] | **7.00E-12** |
| Klemera-Doubal Biological Age | -0.13 | [-0.15,-0.10] | **1.23E-23** | -0.15 | [-0.20,-0.10] | **1.64E-09** |
| Homeostatic Dysregulation | -0.14 | [-0.17,-0.11] | **2.30E-20** | -0.17 | [-0.21,-0.12] | **1.18E-11** |
| <u>DNA Methylation Measures</u> |  |  |  |  |  |  |
| PhenoAge Clock | - | - | - | -0.10 | [-0.15,-0.05] | **9.75E-05** |
| GrimAge Clock | - | - | - | -0.30 | [-0.35,-0.26] | **2.05E-36** |
| DunedinPoAm | - | - | - | -0.20 | [-0.25,-0.15] | **3.87E-15** |
| **Educational Mobility** |  |  |  |  |  |  |
| <u>Blood Chemistry Measures</u> |  |  |  |  |  |  |
| PhenoAge | -0.09 | [-0.11,-0.06] | **2.02E-09** | -0.08 | [-0.12,-0.03] | **1.42E-03** |
| Klemera-Doubal Biological Age | -0.07 | [-0.09,-0.04] | **2.01E-09** | -0.04 | [-0.08,0.00] | 0.068 |
| Homeostatic Dysregulation | -0.06 | [-0.09,-0.03] | **4.18E-06** | -0.06 | [-0.11,-0.02] | **5.90E-03** |
| <u>DNA Methylation Measures</u> |  |  |  |  |  |  |
| PhenoAge Clock | - | - | - | -0.02 | [-0.06,0.03] | 0.465 |
| GrimAge Clock | - | - | - | -0.13 | [-0.17,-0.09] | **3.05E-09** |
| DunedinPoAm | - | - | - | -0.12 | [-0.17,-0.08] | **1.62E-07** |

**Supplemental Table S3.**
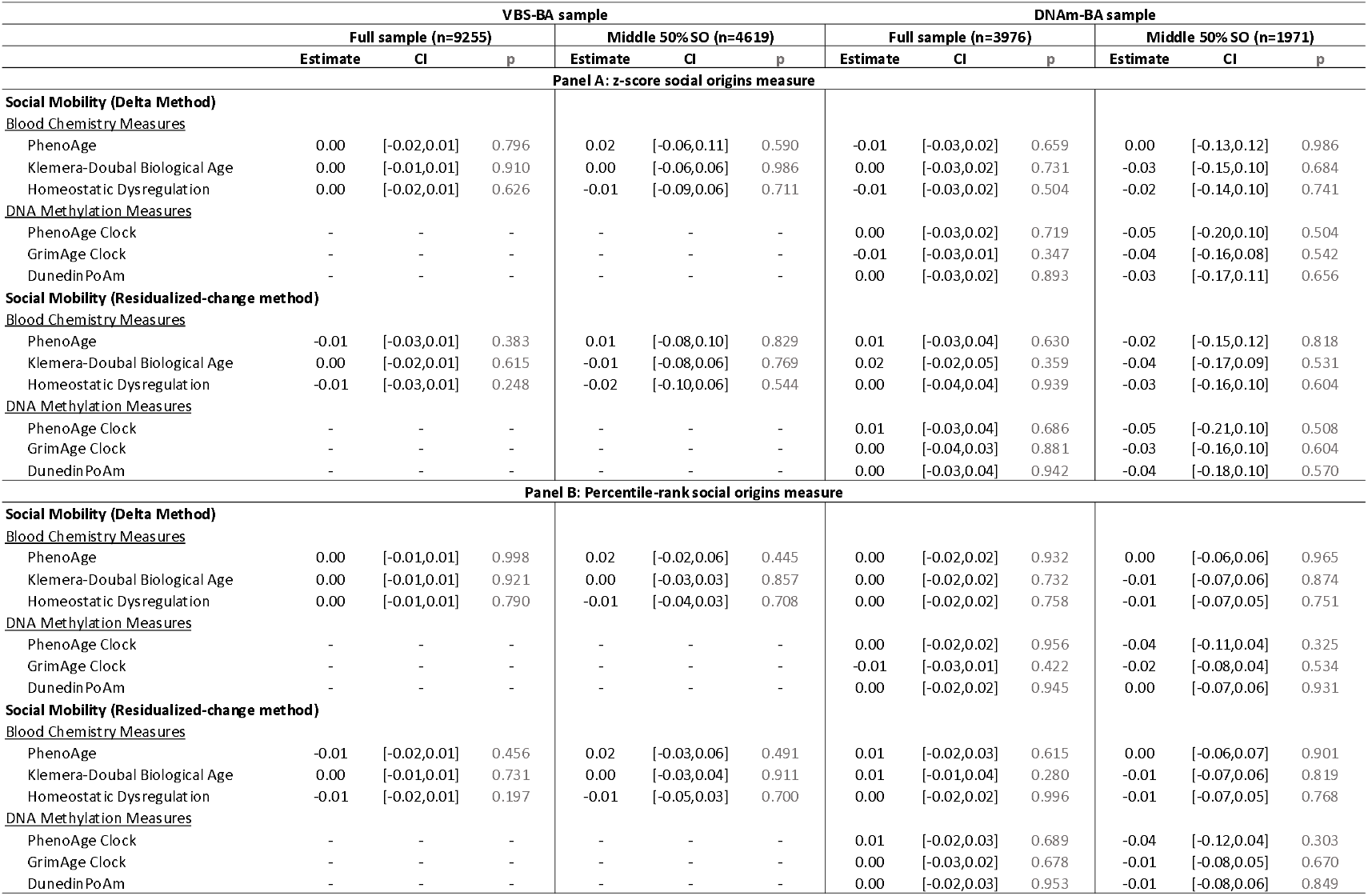
Test of difference in associations of social mobility with biological aging across levels of childhood socioeconomic status. We tested differences in associations of social mobility with biological aging across levels of childhood SES by adding terms to our social mobility regression models for childhood SES level and the interaction of childhood SES level with mobility. The table reports coefficients for interaction terms from these models. Results are presented for the full HRS biomarker sample n=9255) and the HRS DNA methylation sample (n=3786), and for a subset of each sample comprised of participants in the middle 50% of the social origins distribution (HRS biomarker subsample n=4619, HRS DNA methylation subsample n=1890). The purpose of assessing the middle 50% of the social origins distribution was to ensure consistency of results among those whose mobility was not bounded at either end of the distribution.

**Supplemental Table S4a.**
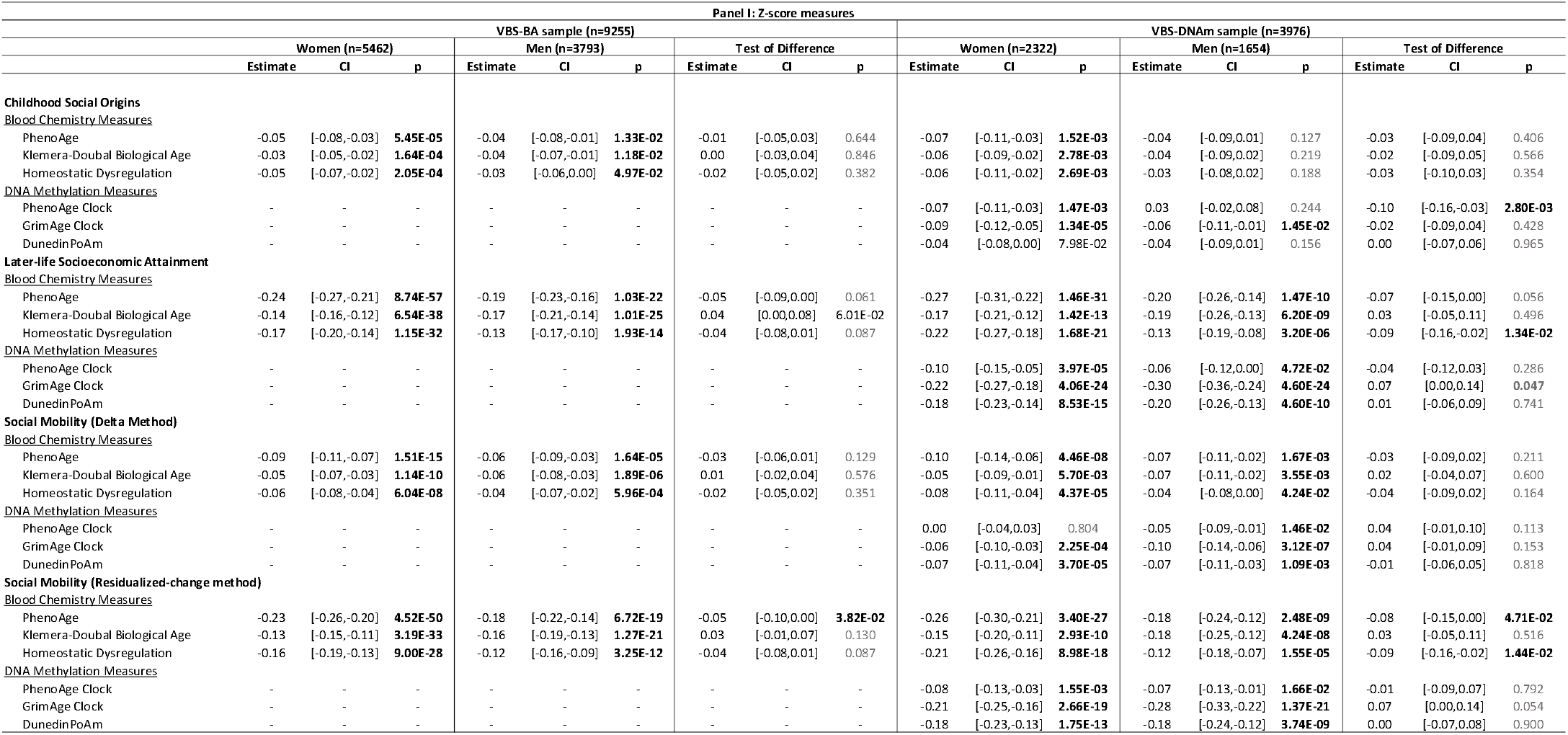
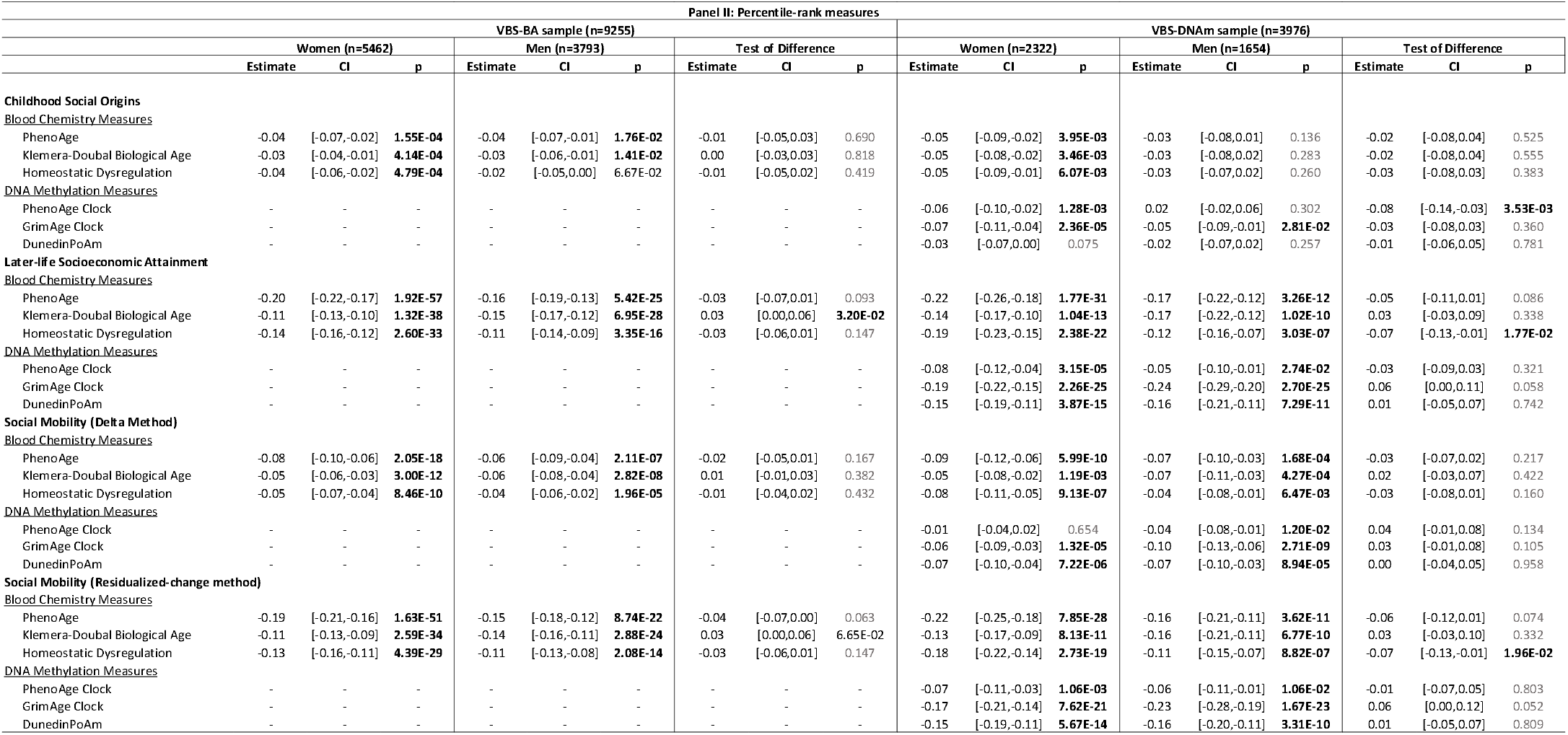
Test of difference in associations of social mobility with biological aging between women and men. We tested differences in associations of social mobility with biological aging between women and men by conducting stratified regression analysis and by adding a term to our social mobility regression models for the interaction of sex with mobility. The first set of columns report results from analysis of women. The second set of columns report results from analysis of men. The third set of columns report the test of difference in results between women and men. This test was conducted by pooling the samples of women and men and fitting the regression model with an additional product term testing the interaction of sex with the measure of social position/mobility. The coefficient reported in this column is the coefficient for the product term testing the interaction. For Z-score measures, effect-sizes are denominated in standard deviation units of biological age advancement per standard-deviation increment in the predictor. For percentile-rank measures, effect-sizes are denominated in standard-deviation units of biological age advancement per 25-percentile-rank increments of the predictor.

**Supplemental Table S4b.**
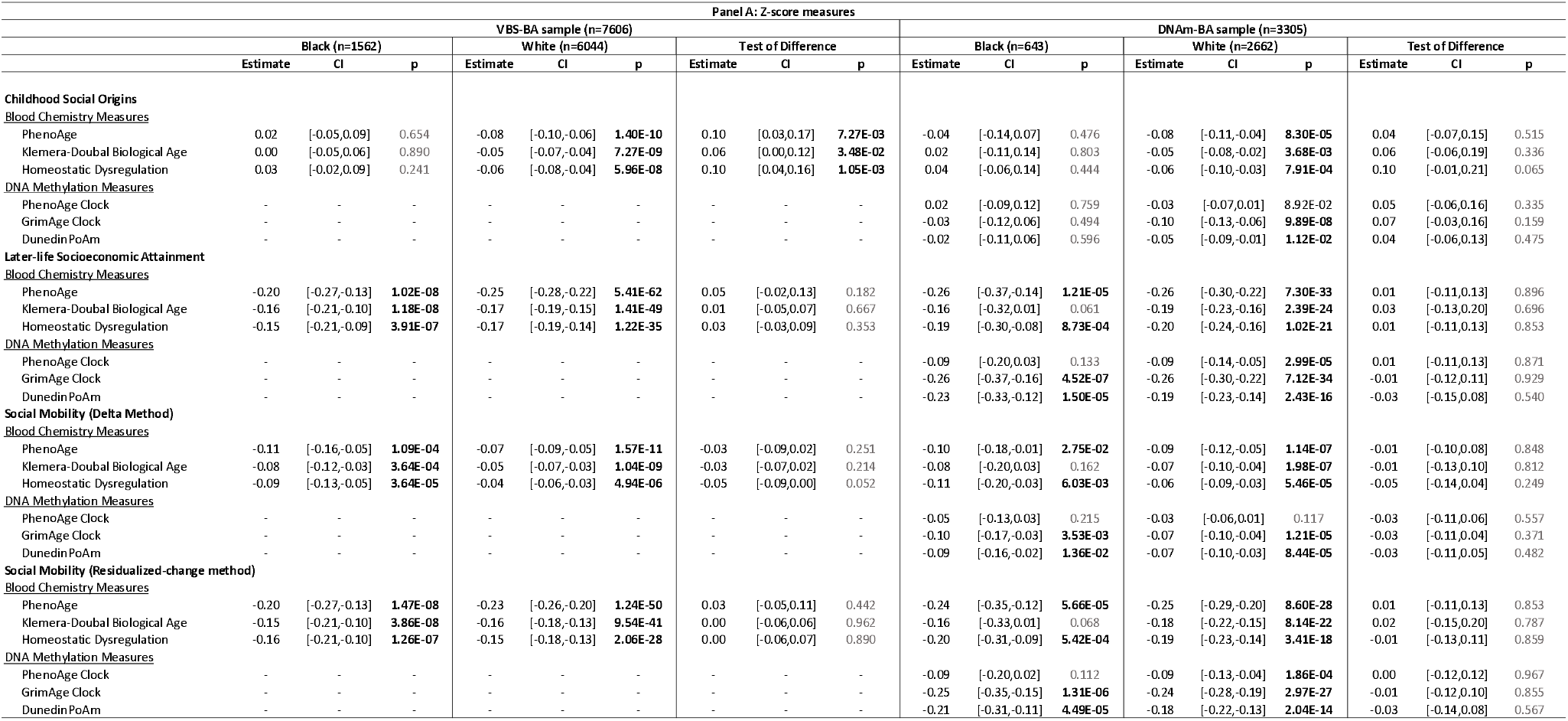
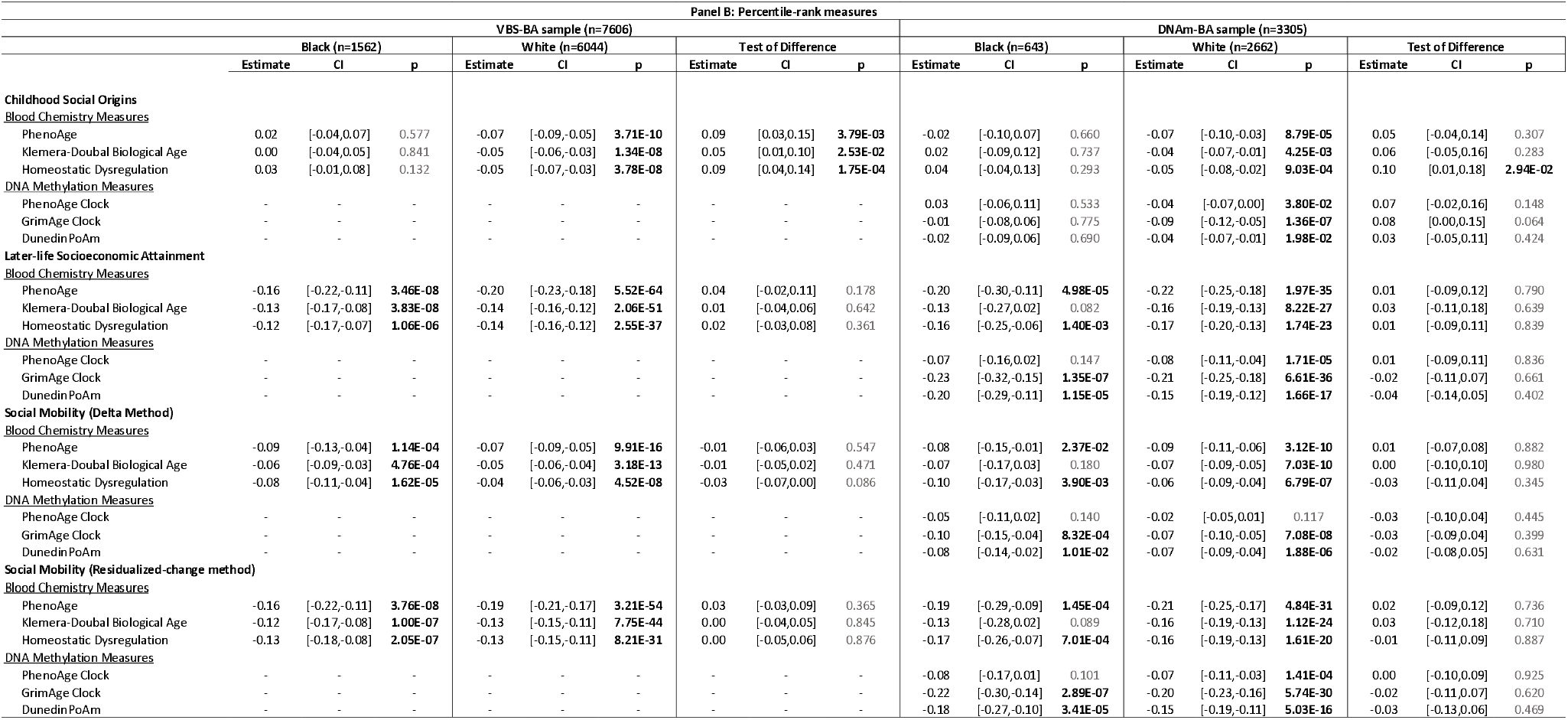
Test of difference in associations of social mobility with biological aging between Black and White participants. We tested differences in associations of social mobility with biological aging between participants identifying as Black and White by conducting stratified regression analysis and by adding a term to our social mobility regression models for the interaction of racial identity with mobility. The first set of columns report results from analysis of women. The second set of columns report results from analysis of men. The third set of columns report the test of difference in results between women and men. This test was conducted by pooling the samples and fitting the regression model with an additional product term testing the interaction of racial identity with the measure of social position/mobility. The coefficient reported in this column is the coefficient for the product term testing the interaction. For Z-score measures, effect-sizes are denominated in standard deviation units of biological age advancement per standard-deviation increment in the predictor. For percentile-rank measures, effect-sizes are denominated in standard-deviation units of biological age advancement per 25-percentile-rank increments of the predictor.

**Supplemental Table S4c.** Test of difference in associations of educational mobility with biological aging between Women and Men. We tested differences in associations of educational mobility with biological aging between participants identifying as Women and Men by conducting stratified regression analysis and by adding a term to our educational mobility regression models for the interaction of sex with mobility. The first set of columns report results from analysis of women. The second set of columns report results from analysis of men. The third set of columns report the test of difference in results between women and men. This test was conducted by pooling the samples and fitting the regression model with an additional product term testing the interaction of sex with the measure of social position/mobility. The coefficient reported in this column is the coefficient for the product term testing the interaction.

|  | VBS-BA sample (n=8759) |  |  |  |  |  |  |  |  | VBS-DNA sample (n=3774) |  |  |  |  |  |  |  |  |
| --- | --- | --- | --- | --- | --- | --- | --- | --- | --- | --- | --- | --- | --- | --- | --- | --- | --- | --- |
|  | Women (n=5193) |  |  | Men (n=3566) |  |  | Test of Difference |  |  | Women (n=2215) |  |  | Men (n=1559) |  |  | Test of Difference |  |  |
|  | Estimate | CI | p | Estimate | CI | p | Estimate | CI | p | Estimate | CI | p | Estimate | CI | p | Estimate | CI | p |
| **Parental education** |  |  |  |  |  |  |  |  |  |  |  |  |  |  |  |  |  |  |
| **Blood Chemistry Measures** |  |  |  |  |  |  |  |  |  |  |  |  |  |  |  |  |  |  |
| PhenoAge | -0.09 | [-0.13,-0.05] | **1.98E-05** | -0.08 | [-0.13,-0.02] | **1.13E-02** | -0.02 | [-0.09,0.06] | 0.654 | -0.09 | [-0.16,-0.03] | **5.38E-03** | -0.11 | [-0.20,-0.03] | **1.11E-02** | 0.02 | [-0.09,0.13] | 0.742 |
| Klemera-Doubal Biological Age | -0.05 | [-0.08,-0.02] | **1.41E-03** | -0.06 | [-0.11,-0.01] | **1.95E-02** | 0.01 | [-0.05,0.07] | 0.697 | -0.12 | [-0.19,-0.05] | **6.26E-04** | -0.13 | [-0.22,-0.04] | **5.52E-03** | 0.01 | [-0.10,0.12] | 0.869 |
| Homeostatic Dysregulation | -0.09 | [-0.13,-0.05] | **2.09E-05** | -0.06 | [-0.11,0.00] | **3.39E-02** | -0.03 | [-0.10,0.03] | 0.347 | -0.11 | [-0.18,-0.05] | **9.38E-04** | -0.10 | [-0.18,-0.01] | **2.55E-02** | -0.02 | [-0.13,0.09] | 0.729 |
| **DNA Methylation Measures** |  |  |  |  |  |  |  |  |  |  |  |  |  |  |  |  |  |  |
| PhenoAge Clock | - | - | - | - | - | - | - | - | - | -0.10 | [-0.17,-0.04] | **2.34E-03** | -0.07 | [-0.16,0.01] | 0.100 | -0.03 | [-0.14,0.08] | 0.561 |
| GrimAge Clock | - | - | - | - | - | - | - | - | - | -0.13 | [-0.19,-0.06] | **1.03E-04** | -0.24 | [-0.32,-0.15] | **4.61E-08** | 0.11 | [0.00,0.22] | **4.14E-02** |
| DunedinPoAm | - | - | - | - | - | - | - | - | - | -0.05 | [-0.12,0.02] | 0.174 | -0.08 | [-0.17,0.00] | 0.063 | 0.04 | [-0.07,0.15] | 0.525 |
| **Participant Education** |  |  |  |  |  |  |  |  |  |  |  |  |  |  |  |  |  |  |
| **Blood Chemistry Measures** |  |  |  |  |  |  |  |  |  |  |  |  |  |  |  |  |  |  |
| PhenoAge | -0.22 | [-0.26,-0.17] | **1.27E-24** | -0.13 | [-0.18,-0.08] | **1.75E-07** | -0.08 | [-0.14,-0.02] | **1.13E-02** | -0.23 | [-0.30,-0.17] | **1.78E-12** | -0.11 | [-0.19,-0.03] | **8.49E-03** | -0.13 | [-0.23,-0.02] | 0.016 |
| Klemera-Doubal Biological Age | -0.12 | [-0.15,-0.10] | **6.79E-18** | -0.13 | [-0.17,-0.09] | **2.72E-09** | 0.00 | [-0.05,0.05] | 0.864 | -0.15 | [-0.21,-0.09] | **5.06E-07** | -0.15 | [-0.24,-0.07] | **4.94E-04** | 0.00 | [-0.11,0.10] | 0.956 |
| Homeostatic Dysregulation | -0.18 | [-0.22,-0.14] | **2.10E-19** | -0.09 | [-0.13,-0.04] | **8.83E-05** | -0.09 | [-0.15,-0.04] | **1.56E-03** | -0.23 | [-0.29,-0.16] | **1.95E-12** | -0.08 | [-0.16,-0.01] | **2.38E-02** | -0.14 | [-0.24,-0.05] | **3.61E-03** |
| **DNA Methylation Measures** |  |  |  |  |  |  |  |  |  |  |  |  |  |  |  |  |  |  |
| PhenoAge Clock | - | - | - | - | - | - | - | - | - | -0.17 | [-0.23,-0.10] | **1.28E-06** | -0.02 | [-0.09,0.06] | 0.618 | -0.15 | [-0.25,-0.05] | **4.10E-03** |
| GrimAge Clock | - | - | - | - | - | - | - | - | - | -0.29 | [-0.35,-0.23] | **3.89E-20** | -0.33 | [-0.40,-0.25] | **2.84E-18** | 0.04 | [-0.05,0.14] | 0.389 |
| DunedinPoAm | - | - | - | - | - | - | - | - | - | -0.19 | [-0.25,-0.12] | **1.47E-08** | -0.22 | [-0.30,-0.14] | **2.97E-08** | 0.04 | [-0.06,0.14] | 0.474 |
| **Educational Mobility** |  |  |  |  |  |  |  |  |  |  |  |  |  |  |  |  |  |  |
| **Blood Chemistry Measures** |  |  |  |  |  |  |  |  |  |  |  |  |  |  |  |  |  |  |
| PhenoAge | -0.11 | [-0.14,-0.07] | **1.06E-08** | -0.06 | [-0.11,-0.02] | **7.36E-03** | -0.04 | [-0.10,0.01] | 0.134 | -0.12 | [-0.18,-0.06] | **6.91E-05** | -0.02 | [-0.09,0.06] | 0.671 | -0.10 | [-0.20,-0.01] | **3.32E-02** |
| Klemera-Doubal Biological Age | -0.07 | [-0.09,-0.04] | **5.25E-07** | -0.07 | [-0.11,-0.03] | **4.90E-04** | 0.00 | [-0.04,0.05] | 0.906 | -0.04 | [-0.09,0.02] | 0.210 | -0.04 | [-0.11,0.02] | 0.203 | 0.01 | [-0.08,0.10] | 0.857 |
| Homeostatic Dysregulation | -0.08 | [-0.11,-0.05] | **5.78E-06** | -0.04 | [-0.07,0.00] | 0.076 | -0.04 | [-0.10,0.01] | 0.098 | -0.10 | [-0.16,-0.04] | **8.26E-04** | -0.01 | [-0.07,0.06] | 0.794 | -0.09 | [-0.18,0.00] | **4.38E-02** |
| **DNA Methylation Measures** |  |  |  |  |  |  |  |  |  |  |  |  |  |  |  |  |  |  |
| PhenoAge Clock | - | - | - | - | - | - | - | - | - | -0.06 | [-0.12,0.01] | 0.082 | 0.03 | [-0.04,0.10] | 0.377 | -0.09 | [-0.18,0.01] | 0.069 |
| GrimAge Clock | - | - | - | - | - | - | - | - | - | -0.14 | [-0.19,-0.08] | **1.72E-06** | -0.12 | [-0.19,-0.06] | **3.59E-04** | -0.01 | [-0.10,0.07] | 0.750 |
| DunedinPoAm | - | - | - | - | - | - | - | - | - | -0.12 | [-0.18,-0.05] | **2.04E-04** | -0.13 | [-0.20,-0.06] | **2.11E-04** | 0.02 | [-0.07,0.11] | 0.695 |

**Supplemental Table S4d.** Test of difference in associations of educational mobility with biological aging between Black and White participants. We tested differences in associations of educational mobility with biological aging between participants identifying as Black and White by conducting stratified regression analysis and by adding a term to our educational mobility regression models for the interaction of racial identity with mobility. The first set of columns report results from analysis of Black participants. The second set of columns report results from analysis of White participants. The third set of columns report the test of difference in results between Black and White participants. This test was conducted by pooling the samples and fitting the regression model with an additional product term testing the interaction of racial identity with the measure of social position/mobility. The coefficient reported in this column is the coefficient for the product term testing the interaction.

|  | VBS-BA sample (n=7257) |  |  |  |  |  |  |  |  | DNA-m-BA sample (n=3161) |  |  |  |  |  |  |  |  |
| --- | --- | --- | --- | --- | --- | --- | --- | --- | --- | --- | --- | --- | --- | --- | --- | --- | --- | --- |
|  | Black (n=1409) |  |  | White (n=5848) |  |  | Test of Difference |  |  | Black (n=587) |  |  | White (n=2574) |  |  | Test of Difference |  |  |
|  | Estimate | CI | p | Estimate | CI | p | Estimate | CI | p | Estimate | CI | p | Estimate | CI | p | Estimate | CI | p |
| **Parental education (composite)** |  |  |  |  |  |  |  |  |  |  |  |  |  |  |  |  |  |  |
| <u>Blood Chemistry Measures</u> |  |  |  |  |  |  |  |  |  |  |  |  |  |  |  |  |  |  |
| Phe noAge | -0.07 | [-0.17, 0.02] | 0.136 | -0.13 | [-0.17, -0.08] | **4.50E-09** | 0.07 | [-0.04, 0.17] | 0.194 | -0.13 | [-0.28, 0.03] | 0.113 | -0.13 | [-0.20, -0.07] | **3.45E-05** | 0.02 | [-0.14, 0.19] | 0.771 |
| Klemera-Doubal Biological Age | -0.04 | [-0.12, 0.03] | 0.272 | -0.08 | [-0.12, -0.05] | **1.64E-07** | 0.04 | [-0.04, 0.13] | 0.279 | -0.22 | [-0.42, -0.01] | **3.55E-02** | -0.09 | [-0.14, -0.04] | **4.26E-04** | -0.12 | [-0.33, 0.08] | 0.245 |
| Homeostatic Dysregulation | -0.05 | [-0.13, 0.03] | 0.194 | -0.10 | [-0.13, -0.06] | **1.48E-06** | 0.07 | [-0.02, 0.15] | 0.134 | -0.11 | [-0.26, 0.03] | 0.131 | -0.10 | [-0.16, -0.04] | **9.95E-04** | 0.00 | [-0.15, 0.16] | 0.967 |
| <u>DNA Methylation Measures</u> |  |  |  |  |  |  |  |  |  |  |  |  |  |  |  |  |  |  |
| Phe noAge Clock | - | - | - | - | - | - | - | - | - | -0.06 | [-0.20, 0.08] | 0.386 | -0.08 | [-0.15, -0.02] | **1.35E-02** | 0.04 | [-0.11, 0.18] | 0.594 |
| GrimAge Clock | - | - | - | - | - | - | - | - | - | -0.08 | [-0.22, 0.06] | 0.273 | -0.21 | [-0.28, -0.15] | **2.14E-11** | 0.14 | [-0.01, 0.29] | 0.070 |
| Duned in PoAm | - | - | - | - | - | - | - | - | - | 0.05 | [-0.09, 0.20] | 0.460 | -0.12 | [-0.19, -0.05] | **3.86E-04** | 0.19 | [0.04, 0.35] | **1.28E-02** |
| **Participant Education** |  |  |  |  |  |  |  |  |  |  |  |  |  |  |  |  |  |  |
| <u>Blood Chemistry Measures</u> |  |  |  |  |  |  |  |  |  |  |  |  |  |  |  |  |  |  |
| Phe noAge | -0.07 | [-0.15, 0.02] | 0.110 | -0.23 | [-0.27, -0.19] | **6.38E-30** | 0.17 | [0.08, 0.26] | **3.07E-04** | -0.06 | [-0.21, 0.09] | 0.430 | -0.23 | [-0.29, -0.17] | **2.02E-13** | 0.17 | [0.01, 0.32] | 0.043 |
| Klemera-Doubal Biological Age | -0.05 | [-0.12, 0.01] | 0.090 | -0.17 | [-0.20, -0.14] | **5.10E-28** | 0.12 | [0.05, 0.19] | **7.15E-04** | -0.15 | [-0.34, 0.04] | 0.130 | -0.17 | [-0.22, -0.12] | **2.13E-10** | 0.02 | [-0.17, 0.21] | 0.840 |
| Homeostatic Dysregulation | -0.09 | [-0.16, -0.03] | **6.76E-03** | -0.16 | [-0.19, -0.12] | **5.67E-18** | 0.07 | [0.00, 0.15] | 0.061 | -0.16 | [-0.30, -0.02] | **2.13E-02** | -0.16 | [-0.22, -0.11] | **1.45E-08** | 0.01 | [-0.14, 0.15] | 0.933 |
| <u>DNA Methylation Measures</u> |  |  |  |  |  |  |  |  |  |  |  |  |  |  |  |  |  |  |
| Phe noAge Clock | - | - | - | - | - | - | - | - | - | -0.04 | [-0.19, 0.10] | 0.549 | -0.12 | [-0.18, -0.06] | **7.83E-05** | 0.08 | [-0.07, 0.24] | 0.299 |
| GrimAge Clock | - | - | - | - | - | - | - | - | - | -0.17 | [-0.30, -0.05] | **6.32E-03** | -0.38 | [-0.44, -0.32] | **8.08E-38** | 0.21 | [0.07, 0.35] | **2.66E-03** |
| Duned in PoAm | - | - | - | - | - | - | - | - | - | -0.23 | [-0.35, -0.10] | **3.32E-04** | -0.24 | [-0.30, -0.18] | **2.47E-14** | 0.03 | [-0.10, 0.17] | 0.636 |
| **Educational Mobility (Index)** |  |  |  |  |  |  |  |  |  |  |  |  |  |  |  |  |  |  |
| <u>Blood Chemistry Measures</u> |  |  |  |  |  |  |  |  |  |  |  |  |  |  |  |  |  |  |
| Phe noAge | 0.00 | [-0.08, 0.07] | 0.936 | -0.11 | [-0.14, -0.07] | **1.01E-09** | 0.10 | [0.01, 0.18] | **2.23E-02** | 0.04 | [-0.10, 0.17] | 0.571 | -0.10 | [-0.16, -0.04] | **3.93E-04** | 0.13 | [-0.02, 0.27] | 0.093 |
| Klemera-Doubal Biological Age | -0.01 | [-0.07, 0.05] | 0.693 | -0.08 | [-0.11, -0.06] | **6.74E-10** | 0.07 | [0.01, 0.14] | **3.38E-02** | 0.04 | [-0.09, 0.17] | 0.572 | -0.08 | [-0.12, -0.03] | **5.62E-04** | 0.12 | [-0.03, 0.26] | 0.107 |
| Homeostatic Dysregulation | -0.04 | [-0.10, 0.03] | 0.256 | -0.07 | [-0.10, -0.04] | **4.01E-05** | 0.02 | [-0.05, 0.09] | 0.504 | -0.04 | [-0.15, 0.08] | 0.507 | -0.07 | [-0.12, -0.02] | **1.04E-02** | 0.02 | [-0.11, 0.15] | 0.754 |
| <u>DNA Methylation Measures</u> |  |  |  |  |  |  |  |  |  |  |  |  |  |  |  |  |  |  |
| Phe noAge Clock | - | - | - | - | - | - | - | - | - | 0.01 | [-0.11, 0.13] | 0.889 | -0.05 | [-0.10, 0.01] | 0.108 | 0.04 | [-0.09, 0.18] | 0.509 |
| GrimAge Clock | - | - | - | - | - | - | - | - | - | -0.07 | [-0.18, 0.03] | 0.183 | -0.17 | [-0.23, -0.12] | **7.56E-10** | 0.10 | [-0.02, 0.22] | 0.111 |
| Duned in PoAm | - | - | - | - | - | - | - | - | - | -0.20 | [-0.31, -0.09] | **5.37E-04** | -0.12 | [-0.18, -0.06] | **3.97E-05** | -0.08 | [-0.21, 0.05] | 0.208 |

**Supplemental Figure S1.**
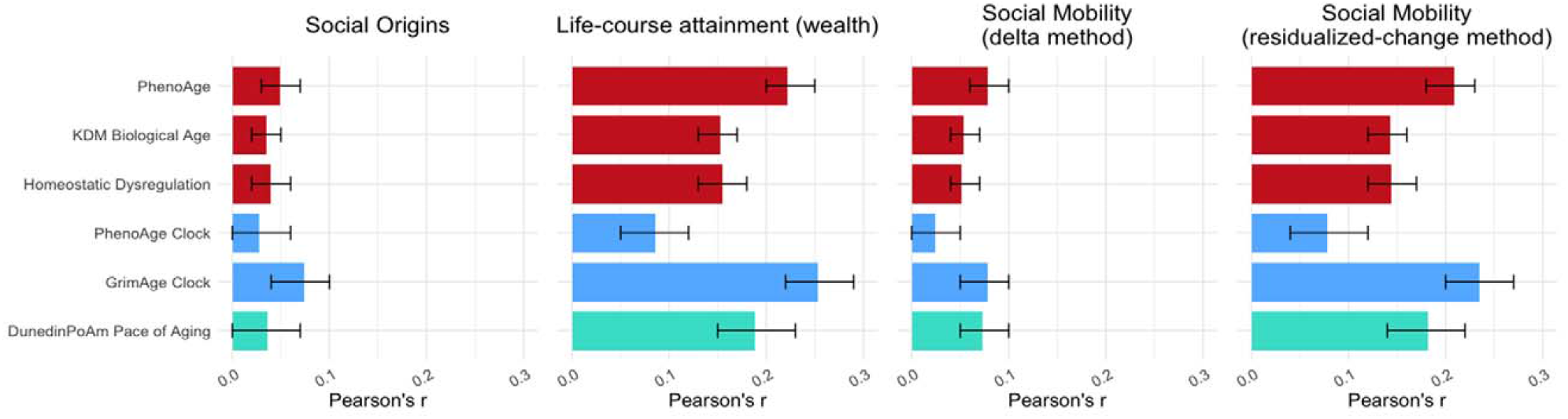
Effect-sizes for associations of childhood and later-life socioeconomic circumstances and social mobility with six blood-chemistry and DNA-methylation measures of biological aging. The figure plots effect-sizes and 95% confidence intervals from analysis of association between measures of social origins, social attainment, and social mobility with six measures of biological aging. Effect-sizes are reported in standard deviation units of the aging measures per standard deviation increment in the predictor, interpretable as Pearson’s r. Blood-chemistry measures are shown in red (n=9255). 2nd generation DNA methylation clocks are shown in blue (n=3976). DunedinPoAm Pace of Aging is shown in turquoise (n=3976). All models are adjusted for age, sex, race, and Hispanic ethnicity.

**Supplemental Figure S2.**
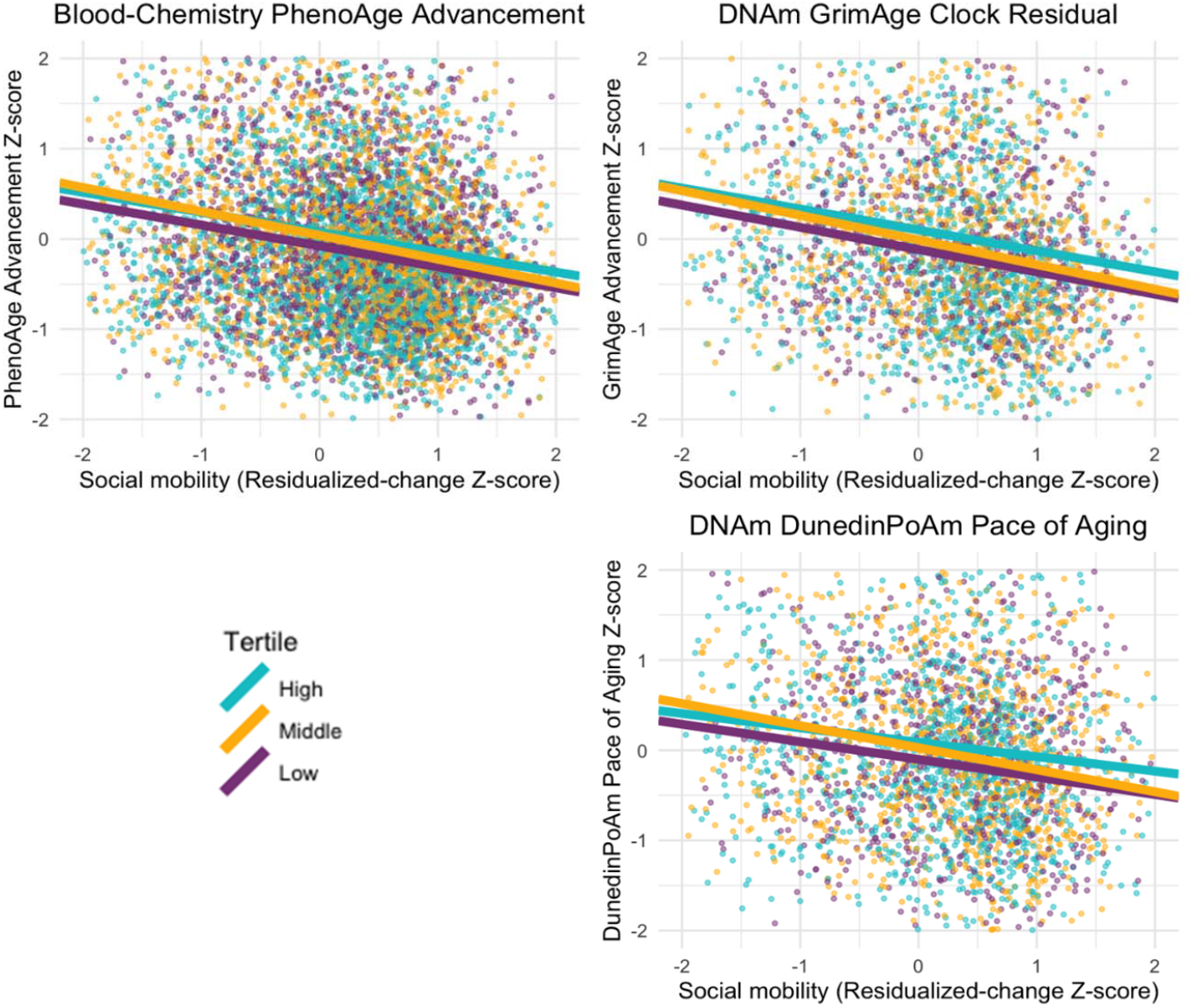
Associations of social mobility with blood-chemistry PhenoAge and DNA-methylation GrimAge and DunedinPoAm by social origins. The figure plots associations social mobility with selected biological aging measures (one blood chemistry clock, one DNA methylation clock, and one Pace of Aging measure). Data are plotted separately for participants who grew up in families with low (purple), middle (orange), and high (teal) socioeconomic status (SES).

**Supplemental Figure S3.**
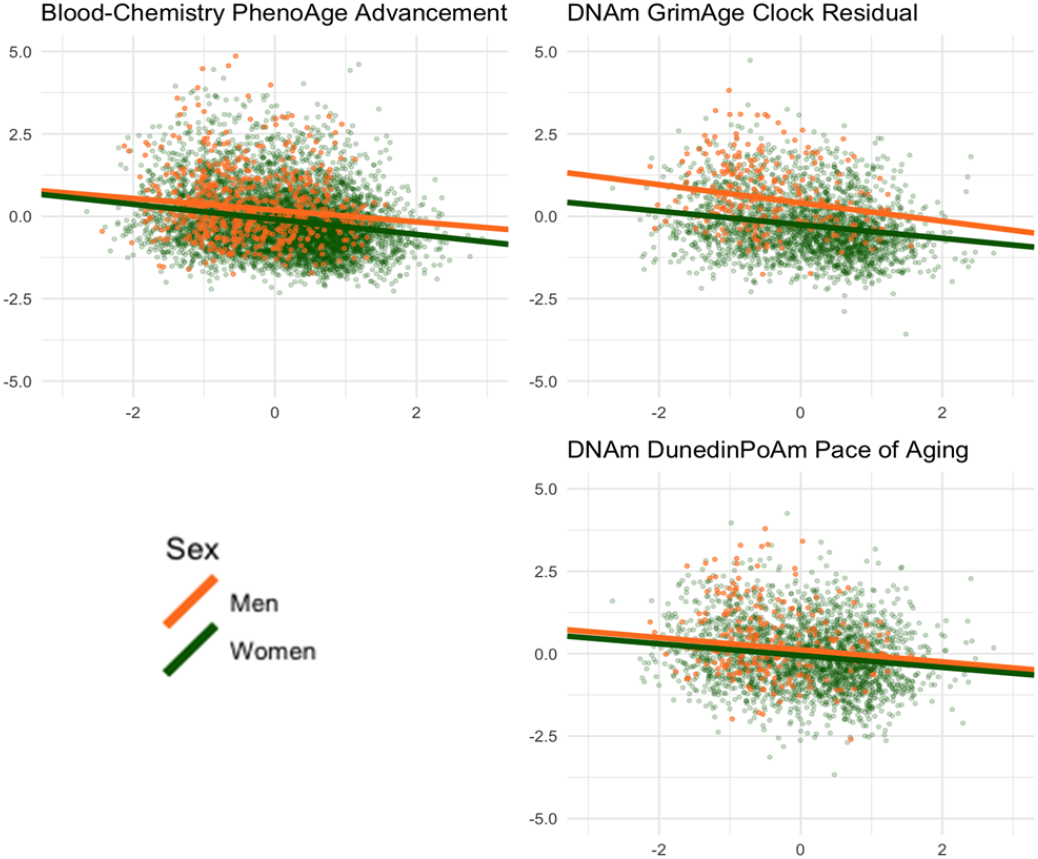
Associations of social mobility with blood-chemistry PhenoAge and DNA-methylation GrimAge and DunedinPoAm in men and women. The figure plots associations social mobility with selected biological aging measures (one blood chemistry clock, one DNA methylation clock, and one Pace of Aging measure). Data are plotted separately for men (orange) and women (green).

**Supplemental Figure S4.**
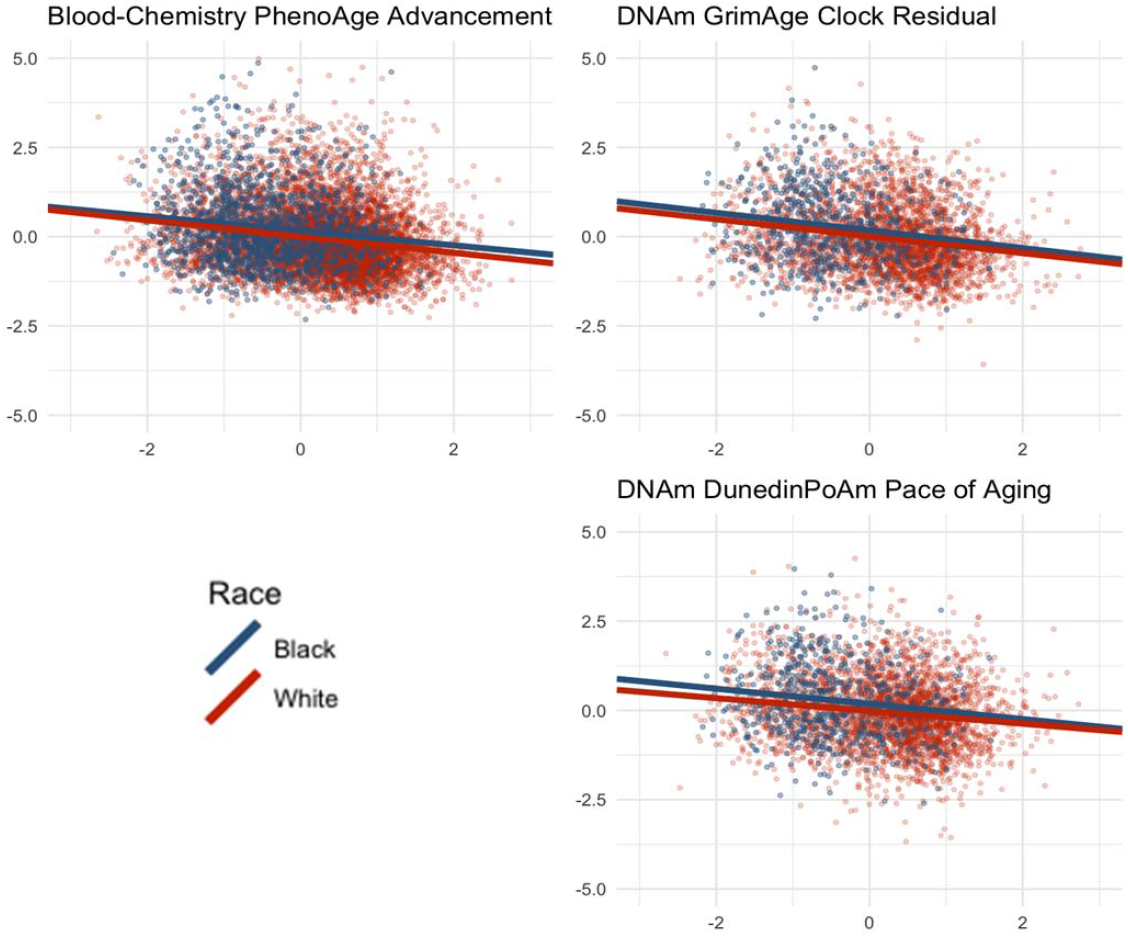
Associations of social mobility with blood-chemistry PhenoAge and DNA-methylation GrimAge and DunedinPoAm in Black and White participants. The figure plots associations social mobility with selected biological aging measures (one blood chemistry clock, one DNA methylation clock, and one Pace of Aging measure). Data are plotted separately for participants reporting Black (blue) and White (red) racial identity.

**Supplemental Figure S5.**
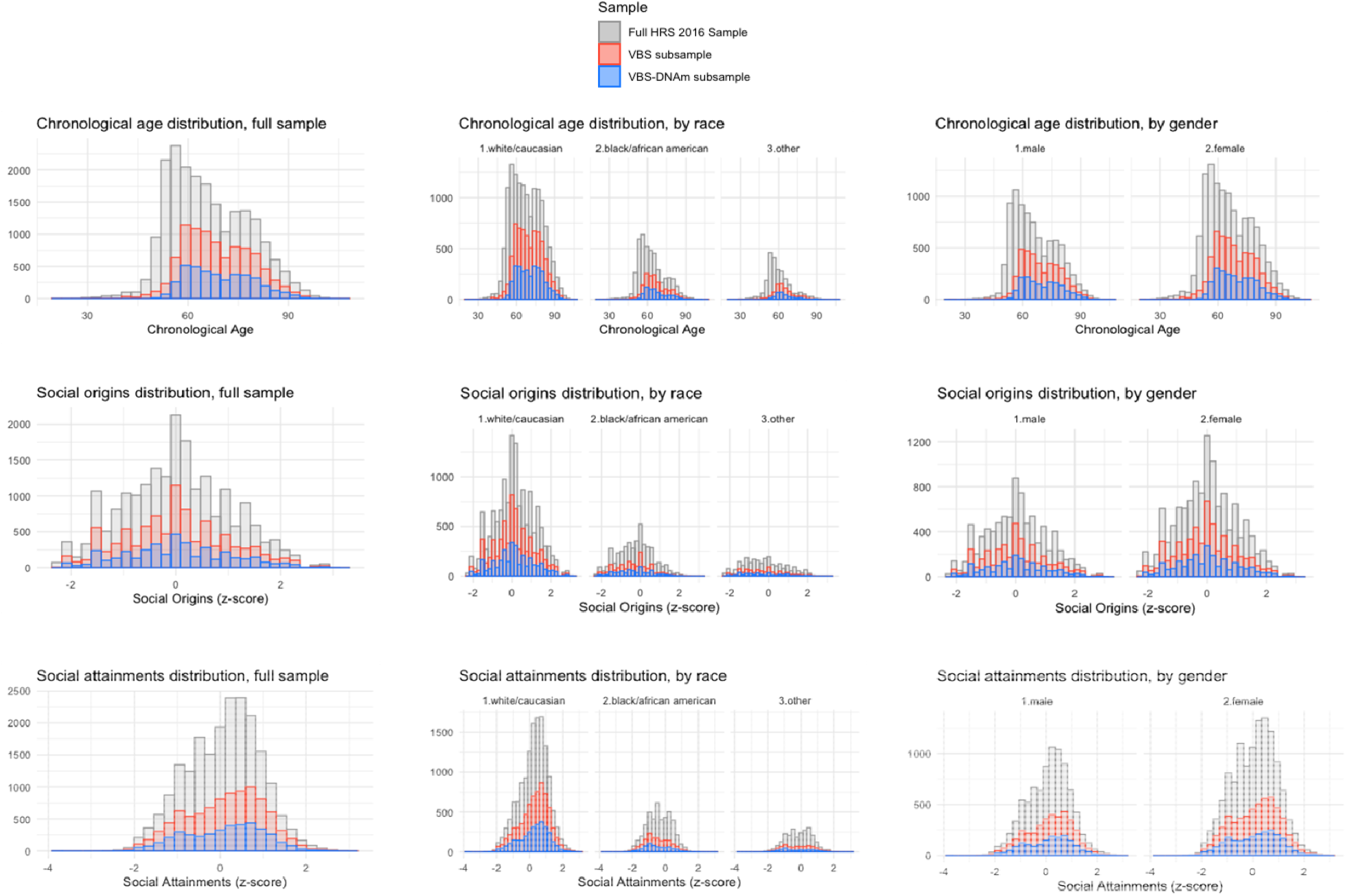
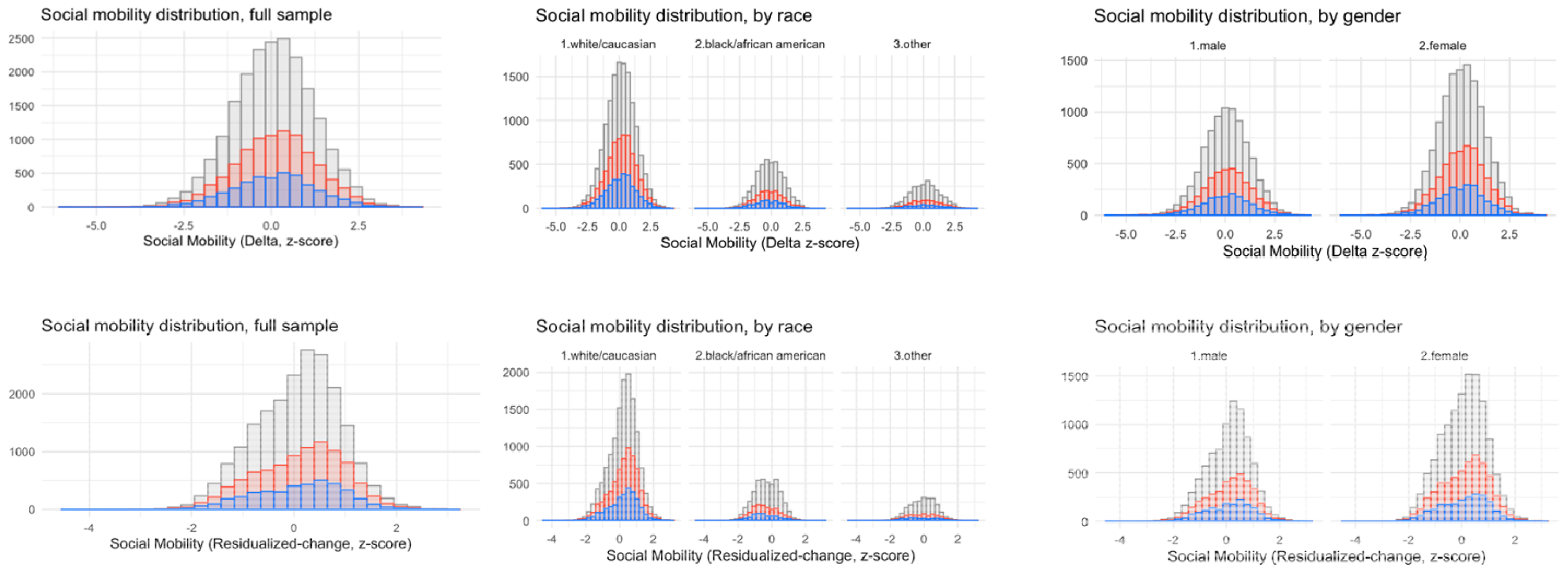
Distribution of demographic and social mobility variables in the US Health and Retirement Study Venous Blood Study and its DNA methylation subsample.

## References

1. N. Snyder-Mackler et al., Social determinants of health and survival in humans and other animals. Science 368 (2020).

2. G. Turrell, J. W. Lynch, C. Leite, T. Raghunathan, G. A. Kaplan, Socioeconomic disadvantage in childhood and across the life course and all-cause mortality and physical function in adulthood: evidence from the Alameda County Study. Journal of Epidemiology & Community Health 61, 723–730 (2007).

3. P. Vineis, M. Kelly-Irving, S. Rappaport, S. Stringhini, The biological embedding of social differences in ageing trajectories. J Epidemiol Community Health 70, 111–113 (2016).

4. J. Campisi et al., From discoveries in ageing research to therapeutics for healthy ageing. Nature 571, 183–192 (2019).

5. L. Ferrucci et al., Measuring biological aging in humans: A quest. Aging Cell 19, e13080 (2020).

6. D. W. Belsky et al., Quantification of the pace of biological aging in humans through a blood test, the DunedinPoAm DNA methylation algorithm. Elife 9 (2020).

7. D. W. Belsky et al., Impact of early personal-history characteristics on the Pace of Aging: implications for clinical trials of therapies to slow aging and extend healthspan. Aging Cell 16, 644–651 (2017).

8. A. Hughes et al., Socioeconomic position and DNA methylation age acceleration across the life course. American journal of epidemiology 187, 2346–2354 (2018).

9. C. McCrory et al., GrimAge outperforms other epigenetic clocks in the prediction of age-related clinical phenotypes and all-cause mortality. The Journals of Gerontology: Series A 76, 741–749 (2021).

10. A. George, R. Hardy, J. C. Fernandez, Y. Kelly, J. Maddock, Life course socioeconomic position and DNA methylation age acceleration in mid-life. J Epidemiol Community Health (2021).

11. S. L. Syme, Psychosocial determinants of hypertension. Hypertension, Determinants, Complications and Intermention, Grune and Stratton, New York, NY, 95–98 (1979).

12. S. A. James, John Henryism and the health of African-Americans. (1994).

13. G. H. Brody, T. Yu, E. Chen, G. E. Miller, Persistence of skin-deep resilience in African American adults. Health Psychology (2020).

14. E. Chen, G. E. Miller, G. H. Brody, M. Lei, Neighborhood poverty, college attendance, and diverging profiles of substance use and allostatic load in rural African American youth. Clinical Psychological Science 3, 675–685 (2015).

15. L. Gaydosh, K. M. Schorpp, E. Chen, G. E. Miller, K. M. Harris, College completion predicts lower depression but higher metabolic syndrome among disadvantaged minorities in young adulthood. Proceedings of the National Academy of Sciences 115, 109–114 (2018).

16. E. M. Crimmins, B. Thyagarajan, M. E. Levine, D. R. Weir, J. Faul, Associations of Age, Sex, Race/Ethnicity and Education with 13 Epigenetic Clocks in a Nationally Representative US Sample: The Health and Retirement Study. The Journals of Gerontology: Series A (2021).

17. E. M. Crimmins, B. Thyagarajan, J. K. Kim, D. Weir, J. Faul, Quest for a summary measure of biological age: the health and retirement study. GeroScience 43, 395–408 (2021).

18. T. B. Kirkwood, Understanding the odd science of aging. Cell 120, 437–447 (2005).

19. Z. Liu et al., Associations of genetics, behaviors, and life course circumstances with a novel aging and healthspan measure: Evidence from the Health and Retirement Study. PLoS medicine 16, e1002827 (2019).

20. M. E. Levine et al., An epigenetic biomarker of aging for lifespan and healthspan. Aging (Albany NY) 10, 573 (2018).

21. D. W. Belsky et al., Eleven telomere, epigenetic clock, and biomarker-composite quantifications of biological aging: do they measure the same thing? American Journal of Epidemiology 187, 1220–1230 (2018).

22. Z. Liu et al., A new aging measure captures morbidity and mortality risk across diverse subpopulations from NHANES IV: a cohort study. PLoS medicine 15, e1002718 (2018).

23. Q. Li et al., Homeostatic dysregulation proceeds in parallel in multiple physiological systems. Aging cell 14, 1103–1112 (2015).

24. G. Fiorito et al., Socioeconomic position, lifestyle habits and biomarkers of epigenetic aging: a multi-cohort analysis. Aging (Albany NY) 11, 2045 (2019).

25. W. J. Hastings, I. Shalev, D. W. Belsky, Comparability of biological aging measures in the National Health and Nutrition Examination Study, 1999-2002. Psychoneuroendocrinology 106, 171–178 (2019).

26. P. Klemera, S. Doubal, A new approach to the concept and computation of biological age. Mech Ageing Dev 127, 240–248 (2006).

27. A. A. Cohen et al., A novel statistical approach shows evidence for multi-system physiological dysregulation during aging. Mechanisms of ageing and development 134, 110–117 (2013).

28. D. Kwon, D. W. Belsky, A toolkit for quantification of biological age from blood-chemistry and organ-function-test data: BioAge. medRxiv 10.1101/2021.08.28.21262759, 2021.2008.2028.21262759 (2021).

29. A. T. Lu et al., DNA methylation GrimAge strongly predicts lifespan and healthspan. Aging (Albany NY) 11, 303 (2019).

30. P. Demakakos, J. P. Biddulph, M. Bobak, M. G. Marmot, Wealth and mortality at older ages: a prospective cohort study. J Epidemiol Community Health 70, 346–353 (2016).

31. J. C. Henretta, R. T. Campbell, Net worth as an aspect of status. American Journal of Sociology 83, 1204–1223 (1978).

32. A. Steptoe, P. Zaninotto, Lower socioeconomic status and the acceleration of aging: An outcome-wide analysis. Proceedings of the National Academy of Sciences 117, 14911–14917 (2020).

33. Health and Retirement Study, RAND HRS Longitudinal File 2018 (V1) public use dataset. Produced and distributed by the University of Michigan with funding from the National Institute on Aging (grant number NIA U01AG009740). Ann Arbor, M1, 2021.

34. T. Friedline, R. D. Masa, G. A. Chowa, Transforming wealth: Using the inverse hyperbolic sine (IHS) and splines to predict youth’s math achievement. Social science research 49, 264–287 (2015).

35. K. M. Pence, The role of wealth transformations: An application to estimating the effect of tax incentives on saving. The BE Journal of Economic Analysis & Policy 5 (2006).

36. C. G. Colen, A. T. Geronimus, J. Bound, S. A. James, Maternal upward socioeconomic mobility and Black–White disparities in infant birthweight. American Journal of Public Health 96, 2032–2039 (2006).

37. R. Chetty, J. N. Friedman, N. Hendren, M. R. Jones, S. R. Porter (2018) The opportunity atlas: Mapping the childhood roots of social mobility. (National Bureau of Economic Research).

38. T. Meschede, J. Taylor, A. Mann, T. M. Shapiro, “Family Achievements?”: How a College Degree Accumulates Wealth for Whites and Not for Blacks. Â€: How a College Degree Accumulates Wealth for Whites and Not for Blacks, 121–137 (2017).

39. W. R. Emmons, L. R. Ricketts, Unequal degrees of affluence: Racial and ethnic wealth differences across education levels. The Regional Economist (2016).

40. M. Kothari, D. W. Belsky, Aging: Unite to predict. Elife 10, e66223 (2021).

41. G. H.-J. Graf et al., Testing Black-White disparities in biological aging in older adults in the United States: analysis of DNA-methylation and blood-chemistry methods. medRxiv 10.1101/2021.03.02.21252685, 2021.2003.2002.21252685 (2021).

42. L. Raffington et al., Socioeconomic disadvantage and the pace of biological aging in children. Pediatrics 147 (2021).

43. J. Wu, K. S. Dean, Z. Rosen, P. A. Muennig, The cost-effectiveness analysis of nurse-family partnership in the United States. Journal of health care for the poor and underserved 28, 1578–1597 (2017).

44. S. Dynarski, C. Libassi, K. Michelmore, S. Owen (2018) Closing the gap: The effect of a targeted, tuition-free promise on college choices of high-achieving, low-income students. (National Bureau of Economic Research).

45. E. Courtin et al., The Health Effects Of Expanding The Earned Income Tax Credit: Results From New York City: Study examines the health effects of the New York City Paycheck Plus program that increases the Earned Income Tax Credit for low-income Americans without dependent children. Health Affairs 39, 1149–1156 (2020).

46. N. Castells, J. Riccio, Executive Skills Coaching Plus Incentives in a Workforce Program: Introducing the MyGoals Demonstration. MDRC (2020).

